# Intelligence and academic performance: Is it all in your head?

**DOI:** 10.1101/2021.01.23.427928

**Authors:** Katherine L. Bottenhorn, Jessica E. Bartley, Michael C. Riedel, Taylor Salo, Elsa I. Bravo, Rosalie Odean, Alina Nazareth, Robert W. Laird, Erica D. Musser, Shannon M. Pruden, Eric Brewe, Matthew T. Sutherland, Angela R. Laird

**Author notes:** Corresponding Author*: Dr. Angela R. Laird, Professor of Physics, Florida International University, Miami, FL, USA |.

## Abstract

Academic performance relies, in part, on intelligence; however, intelligence quotient (IQ) is limited in predicting academic success. Furthermore, while the search for the biological seat of intelligence predates neuroscience itself, its findings remain conflicting. Here, we assess the interplay between IQ, academic performance, and brain connectivity with behavioral and functional MRI data collected from undergraduate students as they completed an active learning or lecture-based semester-long university physics course. IQ (i.e., full-scale WAIS scores) increased significantly pre-to post-instruction, were associated with physics knowledge and reasoning measures, but were unrelated to overall course grade. IQ was related to brain connectivity during physics-related cognition, but connectivity did not mediate IQ’s association with task performance. These relations depended on students’ sex and instructional environment, providing evidence that physics classroom environment and pedagogy may have a gendered influence on students’ performance. Discussion focuses on opportunities to improve physics reasoning skills for all students.

## Introduction

Intelligence testing has been the subject of substantial scientific inquiry over the past century, especially with respect to academic performance and overall success.^1–4^ These lines of inquiry have a pernicious history, as intelligence testing and IQ have been used to deny educational and employment opportunities and as a foundation of eugenics in the United States and abroad in the 20th century, ^5,6^ though updates to these tests have mitigated bias with respect to racial, ethnic, and gender differences.^7,8^ Both in research and in popular discourse, intelligence is often treated as an inherent, stable, trait-like quality, equated with an intelligence quotient (IQ).^9–13^ While intellectual abilities are moderately heritable,^14^ IQ and other psychometric measures of intellectual ability are also influenced by a number of experiential factors, and the relative influences of genes and environment on IQ change across the lifespan.^15,16^ In general, intelligence is psychometrically assessed via a range of verbal and nonverbal cognitive tests, capturing a general view of ability across domains. One such test is the Wechsler Adult Intelligence Scale (WAIS^17^), which demonstrates moderate stability across adulthood, with increasing stability for shorter intervals and with increasing age.^18–28^ A history of research and popular discourse presupposes that IQ predicts one’s predisposition to academic and life success,^29–33^ though in reality, the picture is much more complicated due to a variety of sociocultural factors.^34–37^ Decades of research suggest education and psychometric intelligence are entwined in a bidirectional relationship, as intellectual ability predicts access to and extent of education, through a variety of socioeconomic factors,^38,39^ while years of education predict modest increases in intellectual abilities,^38,40^ and some educational interventions likely to improve one’s ability to acquire and apply knowledge and skills.^22,41–44^

University students who pursue science, technology, engineering, and mathematics (STEM) disciplines are exposed to a rigorous curriculum designed to transform their problem-solving skills,^45–48^ which engages students’ perceptual and verbal abilities.^49–51^ The fourth edition of the WAIS (WAIS-IV) provides a full-scale measure of intellectual ability (FSIQ) and four component index scores: Processing Speed, Perceptual Reasoning, Working Memory, and Verbal Comprehension.^52^ The WAIS is widely used as an extensively validated and researched clinical tool, but less often applied in educational research, though it may be particularly well-suited for exploring associations between education and skill development. Introductory physics presents a prime opportunity for such study, as a gateway course for STEM majors with a relatively standard curriculum across universities, including instruction on classical Newtonian mechanics and emphasizing the development of quantitative, visuospatial reasoning and problem-solving skills likely captured by WAIS-IV index scores. Unfortunately, female students often perform worse on specific conceptual evaluations in these courses, though not necessarily on overall course grades, due to a host of socioaffective and -cultural factors present in education and physics classrooms, specifically.^53–58^ Female students also constitute a smaller proportion of the student body, compared to their male counterparts^53,56,57,59,60^ and ultimately, such disparities can propagate across courses, leading to higher rates of STEM degrees among male students as compared to female students (64% male vs. 36% female in 2015-2016).^61^ Recently, institutions have sought to improve STEM student success using active learning instructional approaches that yield improved student performance outcomes^62,63^ and impact socioemotional aspects of university education, including self-efficacy, science-related anxiety, and identity.^54,59^ Together, these effects may mitigate existing sex differences in performance,^55,58^ though this is contradicted by some findings.^64^ Altogether, there is a need to better understand sex differences in physics education and potential avenues for mitigating these differences, to ensure all students have the opportunity to succeed.

For as long as we have been trying to understand intelligence, we have been searching for its biological substrates. Recently, neuroscience research has studied the underlying neurobiology of intelligence using neuroimaging techniques such as functional magnetic resonance imaging (fMRI).^65–69^ Task-based fMRI research has focused on understanding how differences in brain activation during cognition differs relates to intelligence, yielding two theories of the neurobiology of intelligence: the parieto-frontal integration theory (P-FIT^70^) and the neural efficiency hypothesis (NEH^71^). The PFIT suggests that interactions between frontal and parietal regions underlie intelligence, while the NEH suggests that higher intelligence is reflected by more “efficient” brain activation during cognitively demanding tasks. An alternative view on “neural efficiency” comes from the application of network science to functional connectivity, to show that more intelligent individuals exhibit greater topological efficiency, which describes ease of information transfer across the brain,^66–68,72^ rather than activation efficiency. Much of this work has focused on functional brain connectivity during the resting state, i.e., in the absence of a task or externally-directed cognition,^73–75^ often referred to as “intrinsic” connectivity,^76,77^ mirroring the notion of intelligence as an inherent trait. However, recent work shows that individual differences in intelligence are better predicted by task-evoked connectivity,^78^ presenting an opportunity to merge these two lines of research to better understand individual differences in the neurobiology of intellectual abilities.

Here, we build on the knowledge that the WAIS measures cognitive abilities that are subject to influence by educational interventions, and leverage the WAIS to study individual differences across student performance in an introductory physics course, and potential roles of sex and pedagogy therein. Then, we build on prior neuroimaging research suggesting task-evoked brain organization can explain individual differences in intellectual abilities, to search for a biological substrate for associations between ability and student performance. To do so, we collected data from undergraduate students enrolled in either a lecture-based or active learning section of an introductory physics course. Both pre- and post-instruction, students completed the WAIS-IV alongside a robust fMRI protocol, including two physics-related tasks (Figure 1) with different demands on cognition. The first task engaged students’ reasoning skills and conceptions about forces at work in the natural world (Figure 1A), based on the Force Concept Inventory (FCI^79^; see Bartley et al.^80^ for detailed FCI task results). In this physics reasoning task, students viewed questions about forces on and the movement of objects, along with answer choices that included the correct (e.g., Newtonian) explanation, and choices that reflect common but incorrect (e.g., non-Newtonian) conceptions about forces and motion. The second task required students to recall concepts and equations taught in an introductory physics course (Figure 1B). In this physics knowledge task, students engage semantic memory to recognize equations or definitions of physics concepts learned in the course from a list of possible answer choices presented. Here, we used these data to, first, assess changes in WAIS-IV scores (both FSIQ and index scores) over the course of the semester, then applied a series of linear regressions to assess associations between post-instruction and pre-to post-instruction changes in WAIS-IV scores and post-instruction student performance (i.e., task accuracy and final course grade). Finally, we assessed associations between WAIS-IV scores, and changes therein, and post-instruction functional brain organization during the two tasks, and the degree to which these associations provide a common neural substrate supporting the role of cognitive abilities in student performance. We hypothesized that, while FSIQ itself is stable, different WAIS-IV index scores are differentially associated with performance on physics-related assessments with different cognitive demands. Further, we hypothesized these differences would be reflected in brain organization during these domain-specific tasks (e.g., physics reasoning and physics knowledge), providing a neurobiological explanation for ability-performance relationships across physics-related cognition. Finally, including sex and pedagogy in these assessments may provide an insight into sex differences in physics education and the potential role of active learning in their mitigation, though we did not expect different brain-IQ relations across active learning and lecture classrooms.

**Figure 1.**
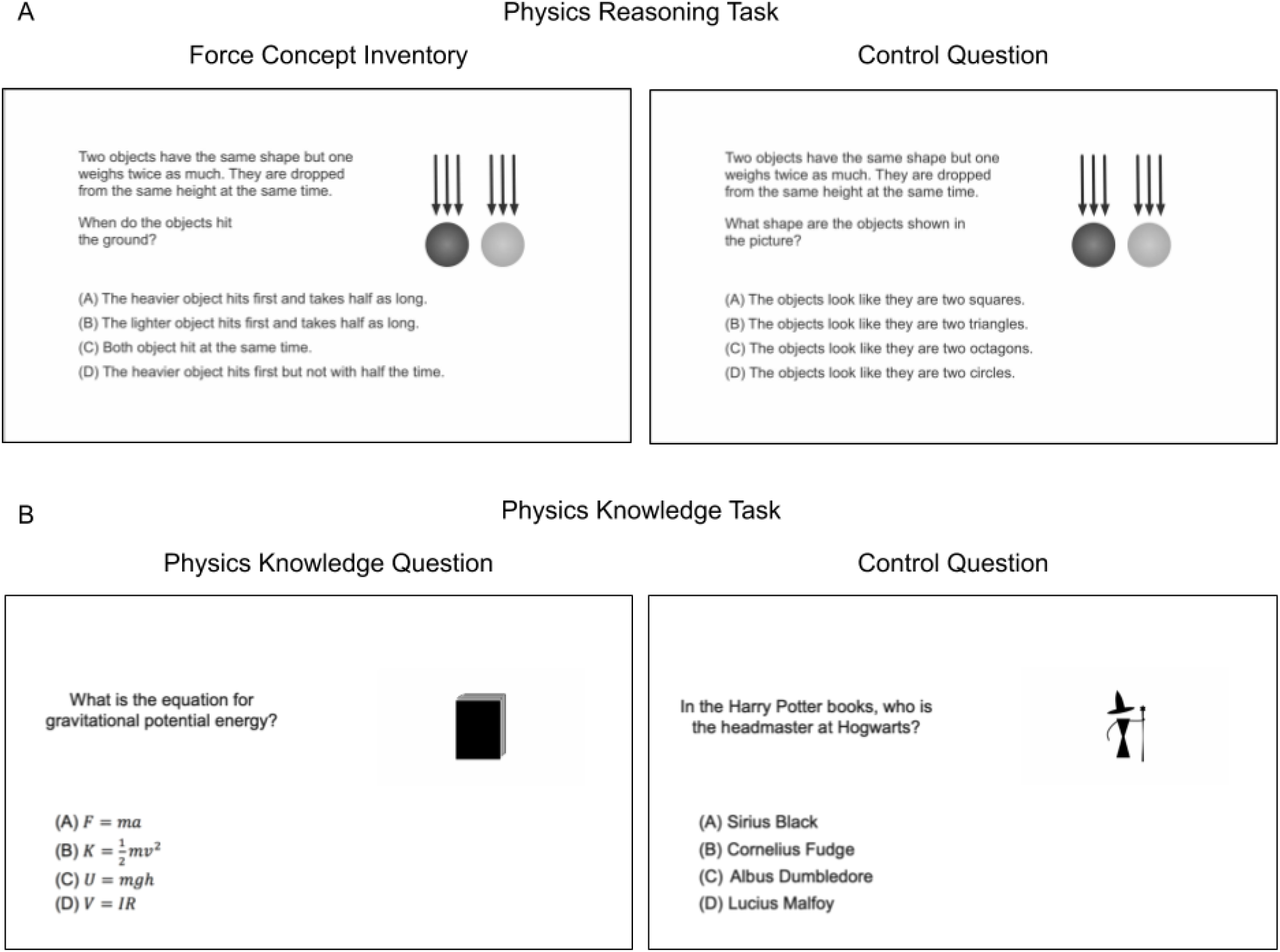
fMRI tasks performed by students at both pre- and post-instruction time points. During MRI scanning sessions, students completed tasks that probed students’ physics-related cognition. The *physics reasoning task* (A) included questions from the Force Concept Inventory (FCI) and engaged students’ conceptual understanding of Newtonian mechanics (left), with perceptually similar control questions (right). The *physics knowledge task* (B) included questions about definitions and equations that students learn in class (left), with perceptually matched general knowledge control questions (right).

## Results

### WAIS-IV scores increased over a semester of physics learning regardless of sex or classroom

First, to understand characteristics of WAIS-IV scores (i.e., FSIQ and index scores) in this sample, we administered the WAIS-IV before and after a semester of physics instruction from 110 students who completed undergraduate introductory physics in lecture-based (25 female, 28 male) or active-learning (24 female, 33 male) classrooms. Pre-to post-instruction changes in WAIS-IV scores were assessed via Wilcoxon signed-rank tests, due to the presence of outliers (Supplementary Figure 1). Full-scale WAIS-IV scores increased an average of 7.03 points (W^+^ = 822, p < 0.001; Table 1; Figure 2A). This change was driven by significant increases in three of the four index scores of the WAIS (Table 1; Figure 2B), greatest in Processing Speed (PSI) and Perceptual Reasoning (PRI). Importantly, however, there were no significant differences in the change in WAIS-IV scores with respect to sex and only PSI changes varied with respect to classroom (i.e., active learning, lecture), evidenced by a significant time by class interaction (Supplementary Table 1). The changes in WAIS-IV scores were commensurate with previously reported retest gains among college students across a similar time period.^81^

**Figure 2.**
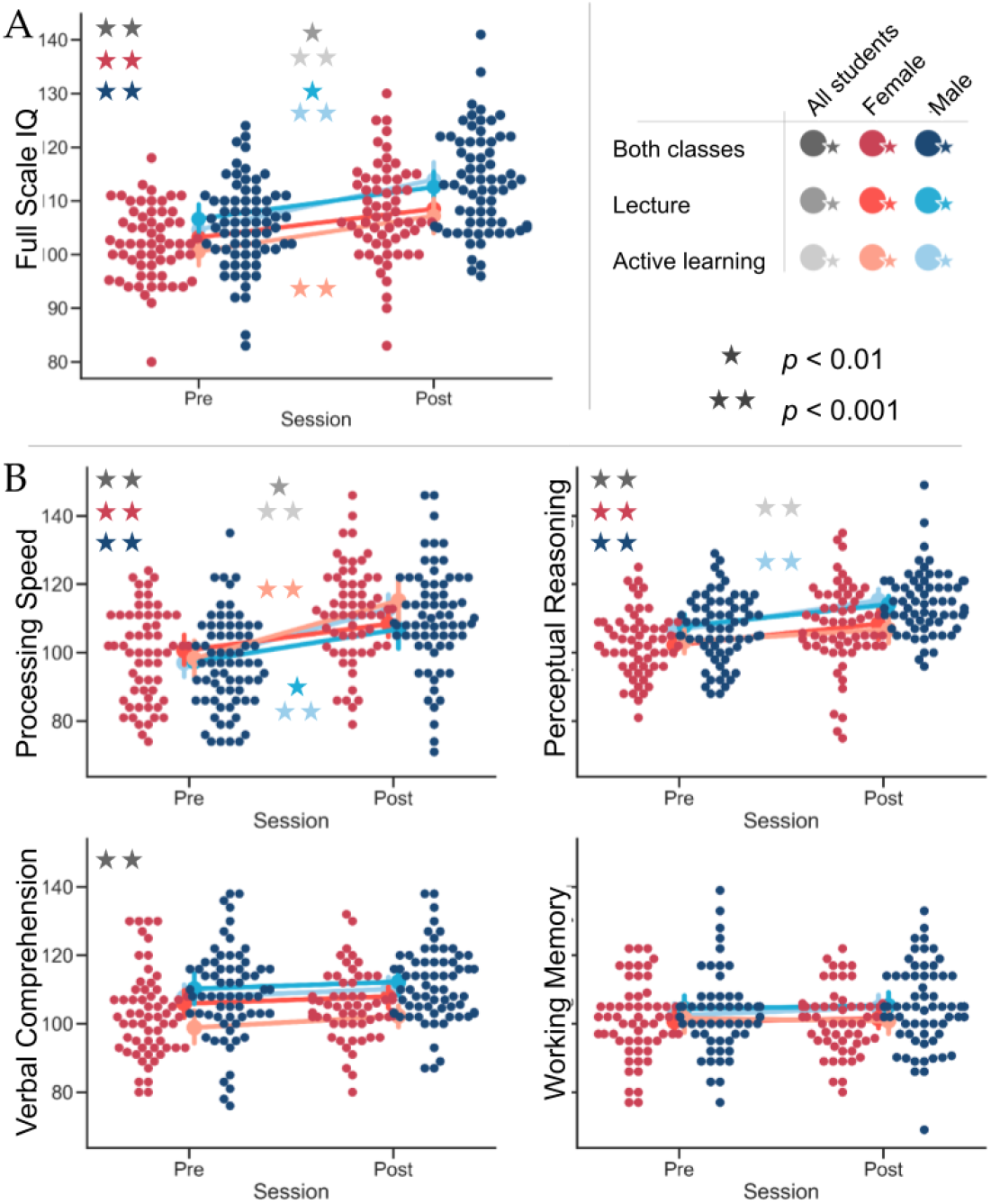
Change in WAIS scores pre-to post-instruction. Full-scale WAIS scores (A) significant increased across students within the sample, for male students in both class types (i.e., active learning and lecture), and for female students in lecture classes. Of the WAIS subscores (B), processing speed (PSI; top left), perceptual reasoning (PRI; top right), and verbal comprehension (VC1; bottom left) showed significant increases across the sample. When separated by sex, only PSI and PRI significantly increased for both sexes, driven by increases in lecture classes. Working memory (bottom right) did not significantly increase for any group in this sample.

**Table 1.** Average change in WAIS scores pre-to post-instruction

| WAIS Score | Pre-instruction | Post-instruction | Pre to Post Change | Estevis et al., 2012 | Change - Estevis *t*-test (*p*) |
| --- | --- | --- | --- | --- | --- |
| Full score<br>WAIS-IV<br>Score | 103.9 ± 7.6 | 110.8 ± 9.4 | **7.0 ± 7.3** | **6.7 ± 5.2** | -0.12 (0.90) |
| Perceptual<br>Reasoning<br>Index | 105.0 ± 9.6 | 111.5 ± 11.1 | **6.0 ± 9.5** | **3.6 ± 5.6** | 2.39 (0.02) |
| Processing<br>Speed Index | 98.3 ± 13.3 | 110.6 ± 14.5 | **12.6 ± 17.5** | **10.5 ± 9.5** | 0.71 (0.48) |
| Verbal<br>Comprehension<br>Index | 106.0 ± 13.0 | 108.5 ± 10.6 | **2.5 ± 9.0** | **4.2 ± 7.6** | -1.28 (0.20) |
| Working<br>Memory<br>Index | 102.3 ± 10.8 | 103.2 ± 10.5 | 2.0 ± 9.1 | 3.2 ± 4.8 | -2.36 (0.02) |

*Note.* Bolded changes indicate significant Wilcoxon Signed Rank tests at $p_{FWE-corr} < 0.05$ ( $\alpha_{adjusted} = 0.014$ ), after controlling for familywise error using the Šidák correction.<sup>81</sup> The *Change - Estevis t-test (p)* column (far right) provides the results of a Student's *t*-test for independent samples, comparing the changes in WAIS scores seen in this sample with those reported by Estevis et al. (2012) to determine whether the changes in our sample are comparable with those reported elsewhere.

### Physics task accuracy, but not course grade, is related to WAIS-IV scores differently for male and female students

To assess associations between course performance and intellectual ability, we separately regressed each WAIS-IV score (both post-instruction and pre-to post-instruction changes) on post-instruction course grade and physics-related task accuracies (denoted “performance” below), controlling for students’ sex, classroom environment, and other demographics, for a total of 30 separate regressions (controlling for familywise error rate with the Šidák correction).

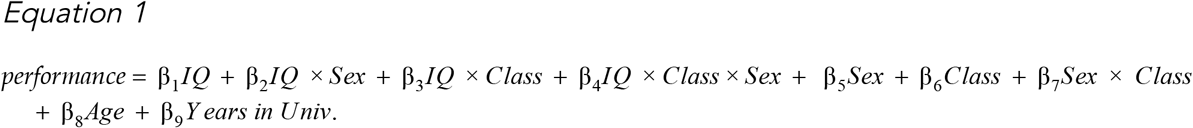

Here, task accuracy refers to the proportion of correct answers given while participants performed a physics task in the MRI scanner (Figure 1). In addition to the full models (Equation 1), we assessed nested models without the WAIS interaction terms (Supplementary Table 2).

Post-instruction accuracy on the physics reasoning task was significantly related to post-instruction PRI scores (post PRI; F(9, 120) = 5.122, p < 0.001; Figure 3A, 3C) and FSIQ scores (post FSIQ; F(9, 120) = 5.770, p < 0.001; Figure 3B), in addition to the pre-to post-instruction changes in both PRI (ΔPRI; F(9, 120) = 5.034, p < 0.001; Figure 3D, 3F) and FSIQ scores (ΔFSIQ; F(9, 120) = 4.498, p < 0.001; Figure 3E, 3G). After controlling for demographics and interactions, post PRI, post FSIQ, ΔPRI, and ΔFSIQ all significantly predicted physics reasoning task accuracy, implying that they were not wholly dependent on students’ sex and classroom environment. However, relations between each post PRI, ΔPRI, and ΔFSIQ and performance were moderated by students’ sex, such that female students exhibited a more positive relationship between WAIS-IV scores and performance than male students (Figure 3B, 3C, and 3D; Table 2, Physics Reasoning Accuracy). Conversely, male students demonstrated overall higher accuracy on the task, in line with previous research.^82^ Of these, the regression of task accuracy on ΔPRI that included class- and sex-interaction terms (i.e., per Equation 1) explained significantly more variance in physics reasoning task accuracy than did the corresponding model without interaction terms. Thus, the relation between task accuracy and ΔPRI is better understood in the context of students’ sex and classroom environments.

**Figure 3.**
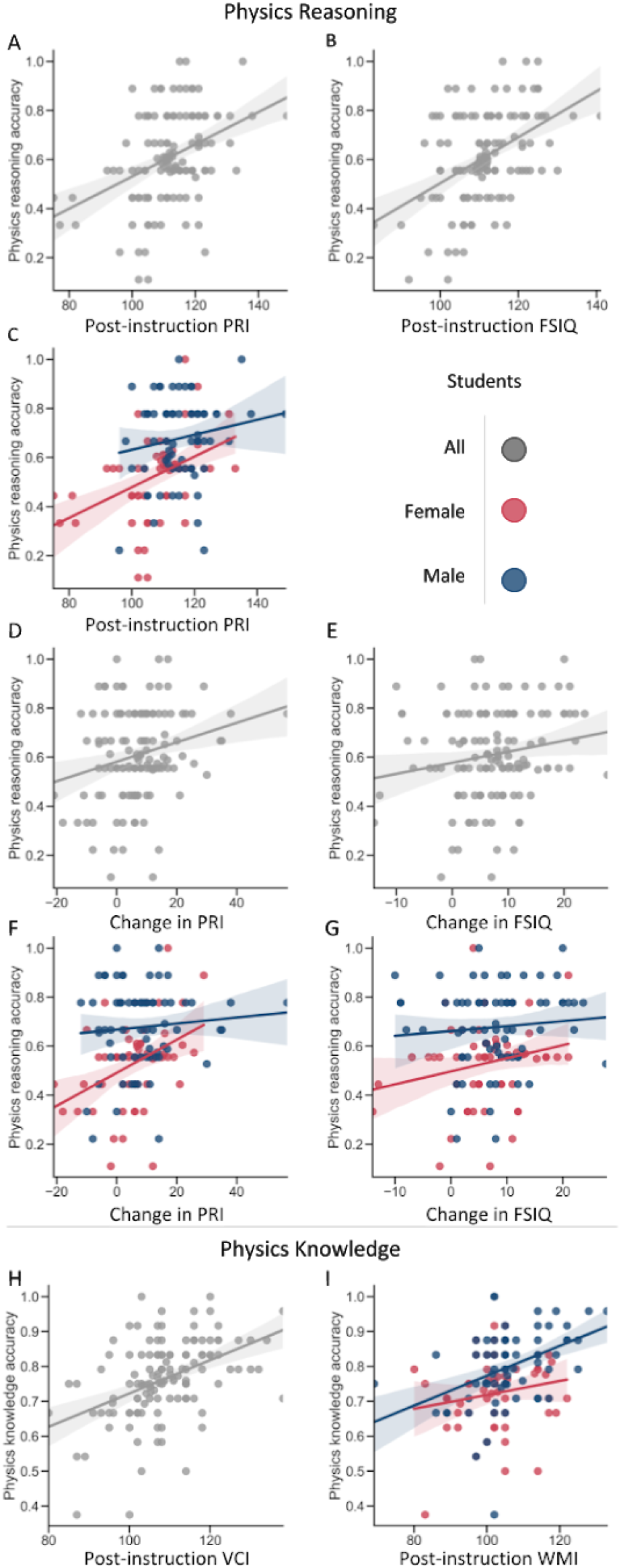
Physics task performance is related to WAIS scores. Post-instruction accuracy on the physics reasoning task was significantly related to post-instruction PRI scores (A) and differently for female and male students (C), and to post-instruction FSIQ scores (B), as well as changes in PRI scores overall (D) and differently for female and male students (F), and changes in FSIQ scores overall (E) and differently for female and male students (G). Accuracy on the physics knowledge task was associated with post-instruction VC1 scores (H) and WMl scores differently for male and female students (I).

**Table 2.** Significant relations between physics task performance and WAIS scores.

|  | *Physics Reasoning Accuracy* |  |  |  | *Physics Knowledge Accuracy* |  |
| --- | --- | --- | --- | --- | --- | --- |
| | Post PRI | Post FSIQ | ΔPRI* | ΔFSIQ | Post VCI | Post WMI |
| WAIS | **0.005** | **0.007** | **0.006** | **0.005** | **0.003** | 0.001 |
| WAIS X Sex (M) | **-0.009** | -0.003 | **-0.010** | **-0.010** | 0.004 | **0.005** |
| WAIS X Class (A) | 0.002 | 0.003 | -0.001 | -0.001 | -0.002 | 0.002 |
| WAIS X Sex X Class | 0.006 | -0.001 | 0.008 | 0.008 | -0.002 | -0.004 |
| Sex X Class | -0.764 | 0.038 | -0.109 | -0.109 | 0.270 | 0.498 |
| Sex (M) | **1.190** | 0.458 | **0.232** | **0.225** | -0.434 | **-0.507** |
| Class (A) | -0.212 | -0.319 | 0.004 | -0.028 | 0.142 | -0.235 |
| Age | **-0.022** | -0.014 | **-0.026** | **-0.029** | -0.001 | -0.004 |
| Year in University | 0.011 | 0.001 | 0.020 | 0.016 | -0.014 | -0.013 |

Regression coefficients are shown for each variable on which physics reasoning and physics knowledge were regressed, for OLS regressions of the form shown in Equation 1 that were significant at $p_{FWE-corr} < 0.05$ and in which a WAIS term or interaction parameter significantly related to performance. Bold text indicates parameters significant at $p < 0.05$ ; bold and italicized text, parameters significant at $p < 0.01$ . \*Indicates that the full interaction model (i.e., of the form shown in Equation 1) explained significantly more variance than a smaller model without interactions between each WAIS scores, class (A for active learning), and sex (M for male).

Post-instruction accuracy on the physics knowledge task was associated with post-instruction Verbal Comprehension Index (post VCI; F(9, 120) = 6.474, p < 0.001; Figure 3H) and Working Memory Index (post WMI; F(9, 120) = 6.008, p < 0.001; Figure 3I). Physics knowledge accuracy was related to post VCI after controlling for potential moderations of this relation by sex and class type (Figure 3H). The converse was true of relations between accuracy and post WMI. Post WMI displayed sex-dependent relations with task accuracy, such that greater accuracy in male students’ was associated with greater increases in WAIS scores than those of female students (Figure 3I). Notably, this moderation was in the opposite direction of those between WAIS scores and physics reasoning accuracy noted above. Of these, regression of task accuracy on WAIS-IV scores with class- and sex-interaction terms (i.e., of the form displayed in Equation 1) did not explain significantly more of the variance in physics knowledge task accuracy than did the models without interaction terms. This indicates that, unlike in the case of physics reasoning, these sex and class interactions do not significantly add to our understanding of these relations.

We found no relations between students’ final course grade and any WAIS-IV score or change therein. Parameters and test statistics for all regressions, including non-significant regressions, can be found in Supplementary Tables 2 and 3.

These data indicate that WAIS-IV scores were clearly, but differentially associated with performance on physics-related assessments, suggesting a distinction between skills related to physics conceptual reasoning and content knowledge recall. Significant associations between physics reasoning accuracy and each ΔPRI and ΔFSIQ suggest that the development of perceptual reasoning ability and general intellectual ability underscore performance in physics reasoning. Similarly, relations between post-instruction VCI and WMI scores and physics knowledge accuracy suggest that working memory and verbal comprehension at post-instruction support successful physics knowledge retrieval, not necessarily the development of those skills (i.e., pre-to post-instruction). Nonetheless, post PRI and full-scale WAIS scores remain relevant for students’ performance on physics conceptual reasoning and problem solving tasks, in addition to the development of such skills.

### Functional brain network efficiency and connectivity differentially support component intelligence across contexts

To further investigate associations between intellectual ability and task performance, we assessed brain organization during physics-related cognition using regressions of the same form as Equation 1 above. Specifically, we combined theories of the neurobiology of intelligence (i.e., P-FIT and NEH) with methods for studying individual differences in the brain organization (i.e., connectomics and network science) to search for a common neural substrate underlying the relations between WAIS-IV scores and accurate physics cognition. Measures of functional connectivity and network efficiency (denoted “topology” below) were regressed on WAIS-IV scores, while students’ sex, classroom environment, demographics, and head movement (i.e., framewise displacement, “fd”).

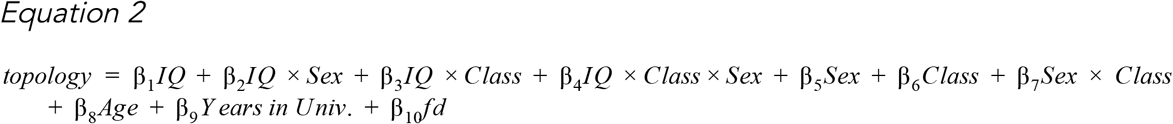

Topological measures were calculated from functional connectivity graphs computed from fMRI data collected while participants performed the physics reasoning and physics knowledge tasks, using two brain parcellations to ensure that results are not parcellation-induced artifacts. In these graphs, individual brain regions comprise nodes and the pairwise correlation of their BOLD signals comprise edge weights, representing functional connectivity. We regressed, separately, (a) global efficiency calculated during each task, (b) local efficiency of each brain region during each task, and (c) connectivity between each pair of brain regions, during each task, on only the WAIS-IV scores significantly related to performance on said task. Significance thresholds of *α* < 0.05 were adjusted to control the familywise error rate using the Šidák procedure, adjusted to account for dependence of correlated measures (Li & Ji, 2005; Sidak, 1967). Regression test statistics, fit statistics, and parameter estimates for all regressions calculated here are shown in Supplementary Tables 4 - 6.

### Global and Local Efficiency

These analyses found that WAIS-IV scores were not significantly associated with global efficiency or with local efficiency across the brain during either task. Across both tasks, only head movement was associated with brain network efficiency.

### Connectivity

Of the WAIS-IV scores associated with physics reasoning task accuracy, only post FSIQ was additionally associated with task connectivity, of the right anterior insula (Figure 4A, 4C), depending on students’ sex and classroom environment (Figure 4B, 4D). For female students greater FSIQ was associated with increased connectivity for those enrolled in lecture-based classes, but decreased connectivity for those enrolled in active learning classes. Meanwhile, for male students, the direction of associations between post-instruction FSIQ and connectivity did not differ due to classroom environment, though it did across parcellations (Figure 4B, 4D). Furthermore, the two parcellations both indicate significant associations of FSIQ and functional connectivity during the physics reasoning task; they did not converge on a particular network or region, providing no specific neuroanatomical locus. Across both parcellations, there was no consistent association between functional connectivity during the physics knowledge task and either post WMI or VCI scores, only with students’ head movement during this task (see Supplementary Figures 2 and 3 for all significant edges).

**Figure 4.**
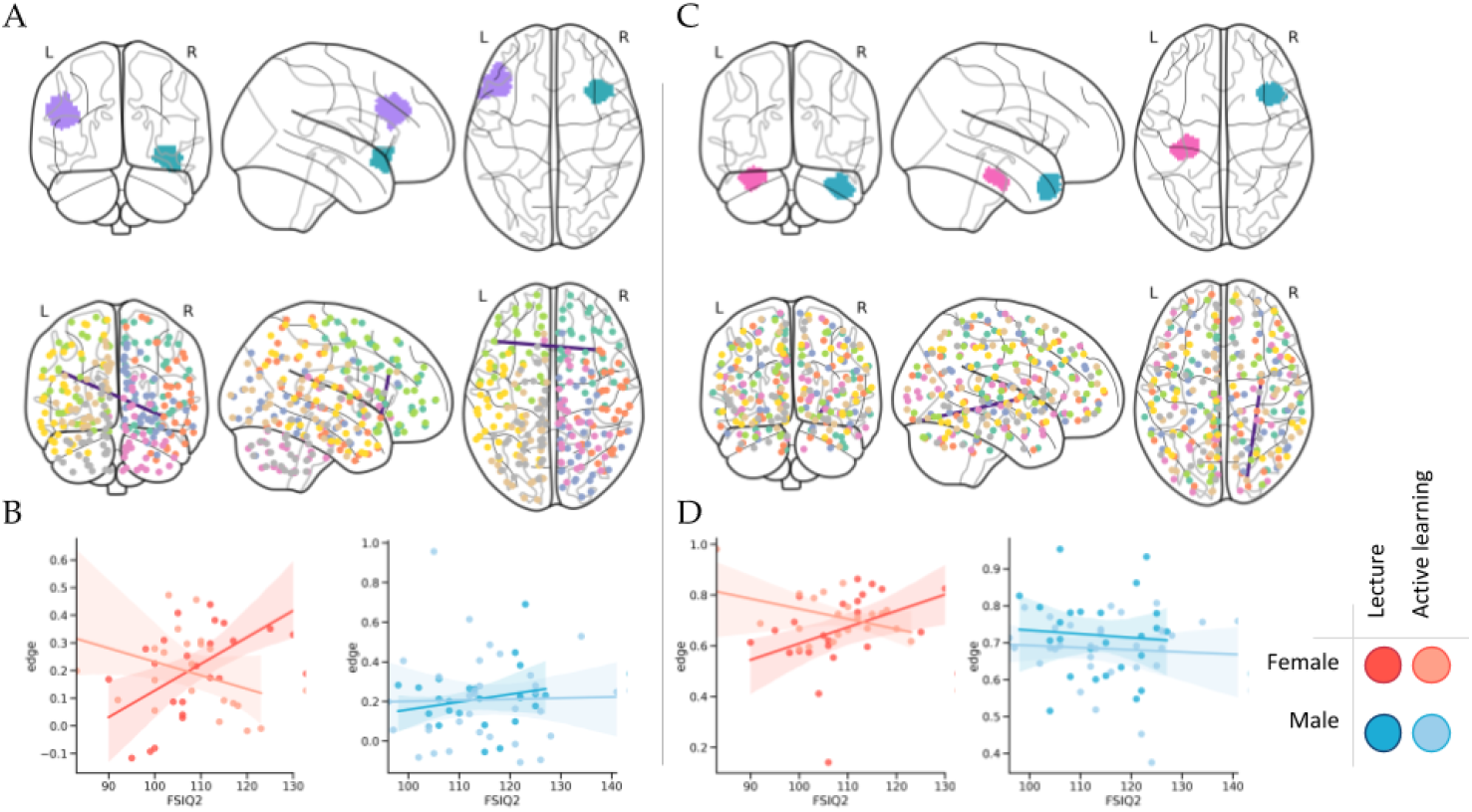
Functional connectivity during physics-related cognition is contextually related to WAIS scores following a semester of physics instruction. Regions (top row) demonstrating functional connectivity (second row) during the physics reasoning task that is significantly related to full-scale IQ scores, color-coded according to anatomical position in the brain to highlight similarities and differences between the parcellations. This connectivity is associated with FSIQ scores differently for male and female students (bottom row) in the Shen (left column) and Craddock (right column) parcellations. Each includes a portion of the anterior insula, though the region to which it is connected differs between parcellations.

### Functional brain networks are not a common neural substrate supporting the role of intelligence in physics-related cognition

We sought to explore possible common neural substrates for WAIS-IV and physics reasoning task accuracy. To this end, we assembled mediation models to assess whether brain connectivity significantly associated with WAIS-IV scores (Table 3, Figure 4) explains shared variance between WAIS-IV scores and task accuracy (Table 2, Figure 3), accounting for the interactions and covariates in Equations 1 and 2 (all model statistics reported in Supplementary Table 7). These models (Figure 5) indicated that the functional connectivity was unrelated to students’ accuracy on the task (yellow path), and while FSIQ scores continued to explain a significant proportion of variability in connectivity (red paths), they no longer significantly accounted for variability in physics reasoning task accuracy (blue paths). Across parcellations, anterior insula connectivity (Figure 4) was significantly associated with FSIQ, but not task accuracy, and provided no significant mediation of the IQ-accuracy relationship (Supplementary Table 3). While WAIS scores and task accuracy seem to capture related behavioral phenomena, FSIQ-accuracy relations were weakened by the inclusion of functional connectivity in the model, and connectivity did not significantly mediate the FSIQ-accuracy association (Table 3).

**Figure 5.**
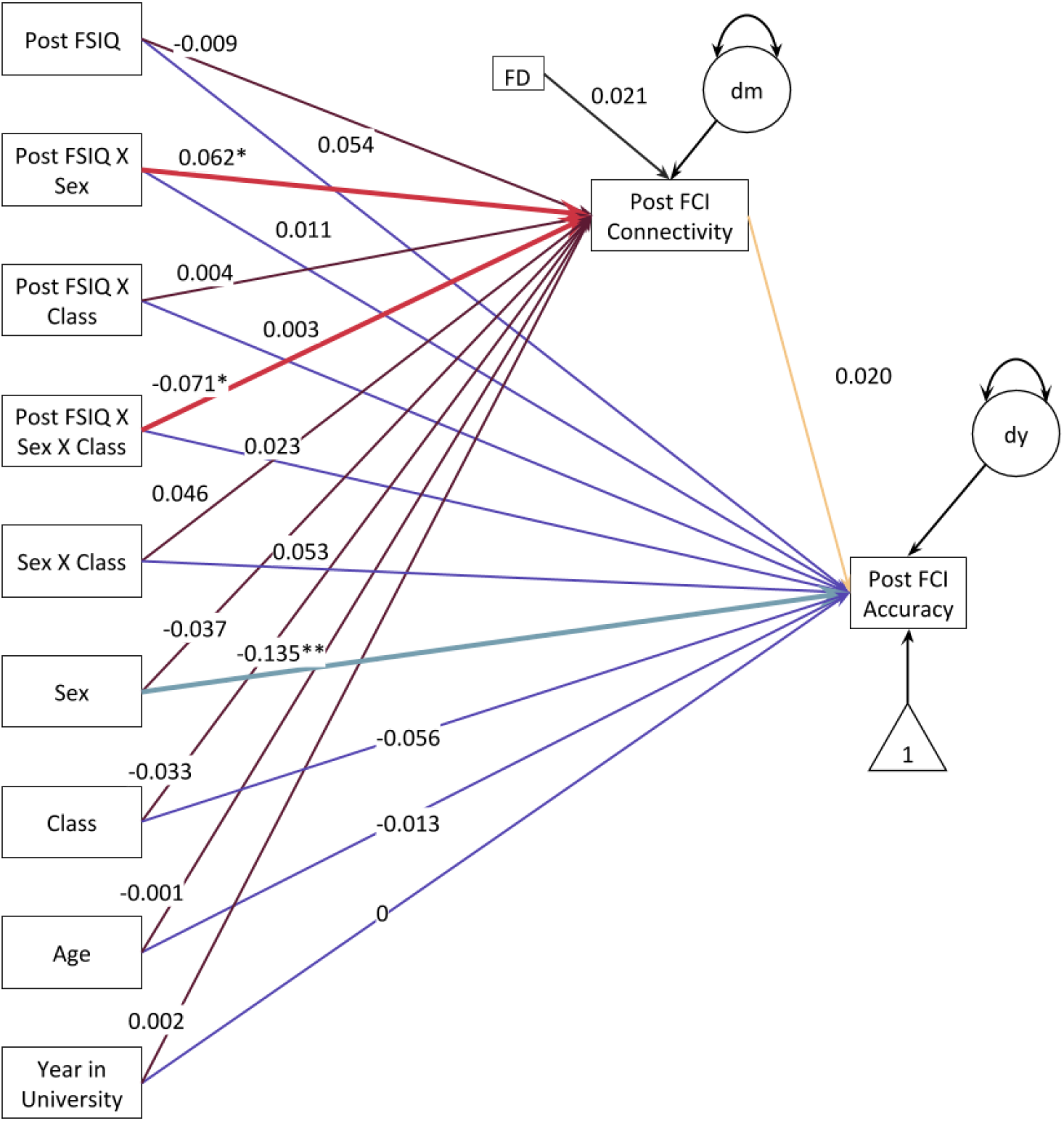
Functional connectivity is related to FSIQ, but does not mediate its relationship with physics reasoning accuracy. This model combines Equations 1 and 2 into a mediation model to assess whether task-based connectivity that is related to full-scale WAIS scores explains the relations between full-scale WAIS scores and students’ accuracy on the physics reasoning task. Overall, this was not the case, although several exogenous variables that significantly explained variance in accuracy (see Equation 1, **Table 2)** were no longer significantly associated with accuracy in this model, where their relations with connectivity are simultaneously being considered, l ighter red paths indicate significant predictors of post-instruction physics reasoning connectivity, while lighter blue paths indicate significant predictors of post-instruction physics reasoning accuracy. *Indicates significance at *p* < 0.05; **, at p < 0.01, model is significant at *p*_FWE-corr_ < 0.01. Covariance between exogenous variables is not shown in this model, but was assessed (see **Supplementary Table** 4).

**Table 3.** Mediation of the relations between changes in full-scale IQ and physics reasoning accuracy by functional connectivity.

|  | Connectivity |  | Accuracy |  |
| --- | --- | --- | --- | --- |
|  | Parameter estimate | P-value | Parameter estimate | P-value |
| FSIQ | -0.009 | 0.695 | 0.054 | 0.123 |
| FSIQ X Sex | **0.062** | 0.025 | 0.011 | 0.805 |
| FSIQ X Class | 0.004 | 0.894 | 0.003 | 0.938 |
| FSIQ X Sex X Class | **-0.071** | 0.043 | 0.023 | 0.691 |
| Sex X Class | 0.046 | 0.199 | 0.053 | 0.355 |
| Sex | -0.037 | 0.26 | **-0.135** | 0.001 |
| Class | -0.033 | 0.154 | -0.056 | 0.191 |
| Age | -0.001 | 0.931 | -0.013 | 0.168 |
| Year in University | 0.002 | 0.467 | 0 | 0.967 |
| Head motion | 0.021 | 0.098 | -- | -- |
| Connectivity | -- | -- | 0.020 | 0.891 |

## Discussion

We present evidence of complex relations between intellectual ability and physics learning, behaviorally and neurally. Among these are significant increases in cognitive ability (i.e., WAIS-IV scores) over a semester of physics instruction, corresponding with previously reported increases in college students tested twice over a similar three-month period. These changes did not differ based on students’ sex or based on course pedagogy and classroom environment. Gains in PRI were among the largest across index scores, and positively related to physics reasoning accuracy, suggesting the WAIS-IV components driving FSIQ increases represent physics-related skill development. Our data indicate that WAIS-IV measures of cognitive ability related to task performance were also related to brain connectivity, but not efficiency, during the task. Although we found no evidence of common neural underpinnings for the performance-ability relationship, we did uncover a moderation of brain-ability associations by students’ sex and physics classroom environment. Therefore, the neurobiology supporting skill acquisition is context-, and perhaps pedagogy-dependent, instead of intrinsic, underscoring the importance of experience and environment in associations between ability and performance.

### Are WAIS gains related to physics education or general college education?

The study of intelligence lacks concrete definitions,^83,84^ but compensates with extensive psychometrics.^38,84–89^ Here, we observed gains in intellectual ability, per the WAIS-IV, that outpaced retest gains reported in the measure’s standardization sample.^17,90,91^ These gains were seen across WAIS-IV index scores, greatest in PSI and PRI though minimal in WMI, but did not significantly differ from retest effects over a similar time period in an independent sample of college students.^92^ WAIS-IV score increases were not related to overall class performance (i.e., course grade), but instead to physics reasoning ability at course completion (i.e., post-instruction FCI accuracy). This suggests they capture skills related to students’ grasp of Newtonian mechanics. While these relations are moderated by students’ class type, we cannot definitively link WAIS-IV score increases to physics instruction, and pedagogical differences therein, without a control group of participants who were not exposed to a semester of physics instruction. Perceptual reasoning seems directly related to skills developed by the specific demands of the course, though it and other components of intelligence are likely developed and engaged broadly across university coursework. To clarify how university instruction may impact different components of intelligence across disciplines, future studies should extend assessments across a range of curricula in the sciences, arts, and humanities and compare retest effects across a semester of education with those seen in the broader population. This is especially relevant given (a) the present evidence of education-related gains in WAIS-IV scores that appear to capture domain-specific skills and, importantly, (b) the widespread belief in the significance of intelligence and IQ for student and life success.

### Perceptual reasoning improvements underlie physics-related cognition

Following a semester of physics instruction, physics knowledge task accuracy was related to post-instruction verbal comprehension and working memory abilities, while physics reasoning task accuracy was related to pre-to post-instruction changes in perceptual reasoning and full-scale WAIS-IV scores (i.e., intellectual or cognitive ability), in addition to their values post-instruction. These differences between the tasks may reflect their unique cognitive demands and the manner in which the associated skills are acquired and exercised.

While physics *knowledge* reflects memorization of formulae and definitions of physics concepts, physics reasoning reflects the development of accurate conceptual understanding of Newtonian mechanics and the macro-scale forces at work in the physical world (e.g., gravity, friction). Students are unlikely to know definitions and formulae learned in class before enrolling in a physics class, but working memory and reading comprehension skills captured by WMI and VCI are not domain-specific but domain-general abilities exercised across curricula.^93–101^ Therefore, it is the associated post-instruction scores that capture the skills utilized in accurately recalling physics knowledge, i.e., recalling definitions and formulae learned in class, that were not necessarily developed during the class.

Conversely, physics *reasoning* draws on the conceptions of physical phenomena that students bring into their first physics class, including pre-existing “common-sense beliefs” acquired over a lifetime of interacting with the physical world that are at odds with scientific explanations.^102^ Decades of physics education research suggests that these pre-existing conceptions are difficult for students to overcome, even with formal instruction.^102–104^ As this instruction relies heavily on visual representations of the movement of macroscale objects to teach Newtonian mechanics, encouraging students to rely on visuospatial skills and mental imagery, likely developing their perceptual reasoning skills throughout the course.^50,105–109^ In this respect, our results indicate that students with more accurate conceptions of Newtonian mechanics following physics instruction were those demonstrating larger increases in perceptual reasoning skills and greater absolute perceptual reasoning ability, post-instruction. It may be that either (a) students who acquire more perceptual reasoning skills in the course are better able to align their conceptions of how the world works with Newtonian explanations or (b) students who are able to overcome previous conceptions about the mechanisms of the physical world gain better perceptual reasoning skills than their peers who have more difficulty doing so. In the former case, these findings might illustrate another approach for physics instructors to help students develop more accurate conceptions of Newtonian mechanics: focusing on students’ perceptual reasoning skills.

Insofar as changes in WAIS-IV index scores reflect skill acquisition, it follows that physics reasoning accuracy, reflecting conceptualizations of Newtonian mechanics honed throughout physics instruction, is associated with *changes* in perceptual reasoning and full-scale intelligence, though *absolute* levels of these skills remain relevant. Conversely, physics knowledge accuracy, reflecting correct recall of definitions and formulae learned throughout physics instruction, is associated with absolute levels of verbal comprehension and working memory skills, but not the changes therein. This may highlight an opportunity for correcting students’ pre-existing conceptions of physical phenomena, by focusing on students’ perceptual reasoning skills throughout their physics instruction. However, the sex- and classroom-differences suggest that this is not a one-size-fits-all opportunity, and that students may benefit differently from such learning interventions. For female students, who historically perform poorer on the Force Concept Inventory (our physics reasoning task) due to a host of sociocognitive factors, increases in PRI might provide a means of “leveling the playing field” compared to their male counterparts.

### Brain network organization during physics cognition is related to intelligence, but does not explain its relationship with physics task accuracy

Contrary to previous research,^73–75^ our data did not indicate associations between students’ WAIS-IV scores and either global or local brain network efficiency and did not support the neural efficiency hypothesis, which suggests more intelligent individuals benefit from “more efficient” brains. The concept of neural “efficiency” is, at best, unclear, with varying definitions across the years, and at worst, empty and misleading.^110^ Here, we use the network science definition of efficiency, defined mathematically in terms of connections between nodes of a graph (i.e., connectivity between brain regions).^111^ These findings add domain-specific insight to what has previously been a study of intrinsic abilities and neurobiological processes, failing to find support in a sample demonstrating intelligence gains related to an extrinsic manipulation (i.e., physics instruction). Our data do not support the parieto-frontal integration hypothesis, either, as we uncovered no parieto-frontal connectivity underlying intelligence during physics-related cognition.

Representing a small proportion of all possible connections in the brain, sparse connectivity during physics reasoning was related to post-instruction full-scale WAIS-IV scores, the full complement of measured verbal, perceptual, working memory, and processing speed skills. Different roles of pedagogy and classroom environment on brain network connectivity, cognitive abilities, and relations between the two with respect to sex point out a potentially significant sex difference in classroom experience. In a heavily male-dominated field like physics, it is a reasonable assumption that male and female students would have differential classroom experiences.^60,112–115^ For example, it is a commonly-held stereotype that men are good at math and that women are not,^116,117^ subjecting women and female students to stereotype threat in physics classrooms, where beliefs negatively affect classroom experience and performance.^118,119^ The data presented here indicate no sex differences in overall course performance, but persistent sex differences across classroom environments in associations between cognitive abilities and not only performance on physics assessments, but brain connectivity during those assessments. Together, this literature and our findings indicate meaningful neurobiological consequences of classroom experience on the basis of sex. However, our data show no sex- or classroom-related difference in changes in intelligence or cognitive abilities, though the brain-WAIS relationships point to a multitude of neurobiological representations of intelligence. Neural phenomena related to intelligence differ based not only on cognitive context, but on sociological and pedagogical contexts, as well.

Although intelligence measures associated with physics task accuracy also explained certain functional connectivity during these tasks, we found no evidence of a common neural substrate for intelligence and accuracy. On the whole, task-based connectivity related to intelligence was unrelated to participants’ accuracy on said task, and no connectivity associated with intelligence accounted for any significant portion of the relations between intelligence and accuracy. This may indicate a lack of a common neural substrate for these two phenomena, suggesting that, while they are measuring a similar skill or capacity, they are not measuring the same skill or capacity. On the other hand, the neural substrate of a common skill measured by PRI or full-scale WAIS scores and physics reasoning accuracy may merely lie beyond a linear relation with brain network connectivity or topology. In any case, understanding the neural instantiation of a common reasoning skill to perceptual reasoning and physics reasoning requires further study.

Finally, rather than a global property of brain network organization, as indicated in prior research,^66,68,120^ these data indicate that sparse, coordinated interactions of disparate brain regions underlie intelligence, in this domain-specific context. That connections across the brain during physics-related cognition are related to changes in students’ overall intellectual skills, but differently with respect to their classroom environment, casts further doubt on the notion of IQ or intelligence as a fixed, innate measure and, instead, highlights the role of environment and experience. Although differences in these relationships between female and male students support the substantial body of literature supporting this notion of sex differences in the biological representations of intelligence,^71,121–128^ here we suggest a potential sociological explanation. These findings indicate, too, neural support for intelligence exhibits domain-specific relations in the context of STEM education. Not only does cognitive context matter to the relations between intelligence and brain network organization, our data indicate that sex and learning environment matter, too.

### Limitations and future directions

Here, we demonstrate increases in intellectual ability over a semester of physics instruction. While students’ sex and classroom environment did not affect the extent of these increases, the data suggest meaningful consequences of classroom environment on the relations between intellectual ability and underlying brain network organization. The implication that the learning environment affects male and female students differently, both cognitively and neurobiologically, in a field as male-dominated as physics demands attention and further study. However, as there were no observed sex differences in final course grade or in change in intellectual ability, any differences in classroom experience are not differentially affecting female and male students’ academic performance in the course. Further work should assess whether differences in experience and associated brain function are linked to long-term success for male and female students. This assessment should consider factors beyond overt measures of success and focus on variables related to self-efficacy and in-classroom experiences, both previously been shown to affect men and women differently in physics education.^58^

While we have identified differences in physics-related brain organization and its relation to WAIS-IV scores based on class type, this study is unable to distinguish whether these differences are due to differences in pedagogy or social classroom environment, or to practice effects across a short assessment period. Future research should include in-classroom assessments of social climate and the possibility of gender differences in social interactions during physics instruction. Furthermore, control groups in (1) another, less male-dominated, domain and (2) an age- and sex-matched groups outside of university would provide insight into both sex differences in and the degree to which changes in intellectual ability are associated with STEM education.

## Conclusion

While our data indicate clear relations between domain-specific components of intellectual ability and performance on assessments of physics conceptual reasoning and content knowledge, we found no association between academic outcomes and intelligence. Likewise, intelligence was related to functional brain connections during these assessments, but none that explain its association with performance. Our multifaceted approach to studying the neurobiological underpinnings of intelligence did not uncover a single, robust aspect of brain network organization that was consistent across cognitive contexts and experiential influences. However, relations between intelligence and, separately, physics task performance and task-based functional connectivity were moderated by students’ sex and classroom environment. Ultimately, both the magnitude and development of perceptual reasoning skills is meaningful over the course of a semester of physics instruction. As these data show significant increases in perceptual reasoning over a semester, we present an optimistic and more plastic view of intelligence, as a set of skills to be developed, rather than an innate capacity that students either have or don’t have. Together, these data highlight the complicated nature of relations between intelligence and classroom successes, which vary with students’ sex and domain-specific experiences.

## Methods

### Participants and Study Design

One hundred and thirty healthy right-handed undergraduate students (mean age = 20.03 ± 2.25 years, range = 18-25 years; 61 females) who completed a semester of introductory calculus-based physics at Florida International University (FIU), a Hispanic Serving Institution, took part in this study. Participants were not currently using psychoactive medications and reported that they had not been diagnosed with any cognitive impairments or neurological or psychiatric conditions. The physics course emphasized problem solving skill development and covered topics in classical Newtonian mechanics, including motion along straight lines and in two and three dimensions, Newton’s laws of motion, work and energy, momentum and collisions, and rotational dynamics. Students were either enrolled in a lecture class or an active learning, “Modeling Instruction”, class, which bases course content in conceptual scientific models and instructs students to appropriate scientific models for their own use. Students completed behavioral assessments and MRI scans in separate appointments at two time points: at the beginning (“pre-instruction”) and conclusion (“post-instruction”) of the 15-week semester. Pre-instruction data collection sessions were acquired no later than the fourth week of classes and post-instruction sessions were completed no more than two weeks after the final exam. Written informed consent was obtained in accordance with FIU’s Institutional Review Board approval.

### Missing Data

A missing value analysis indicated that, of the variables of interest in this study, missingness ranged from 2% to 17%. Data were more often missing from MRI data than behavioral or demographic data and more often missing from post-instruction data than from pre-instruction. Assessment of the relations between missingness on each variable and values of each other variable of interest revealed that data were likely missing completely at random (MCAR). Behavioral and brain network efficiency data were imputed using iterated Bayesian ridge regression implemented in scikit-learn (v. 0.23.1; scikit-learn.org/). Due to its high dimensionality, missingness in edgewise functional connectivity data was addressed using distance-weighted *K*-Nearest Neighbors approach (*K* = 100, where p = 71,824) implemented in scikit-learn, which is robust to missingness up to 20% ^129^.

### Behavioral Measures

During pre- and post-instruction behavior sessions, participants were administered the fourth edition of the Wechsler Adult Intelligence Scale (WAIS-IV^17^), a standardized intelligence test for adults, in addition to other assessments not used here. The WAIS-IV provides scores in four domains, in addition to an overall score of intellectual functioning. The Verbal Comprehension index measures application of verbal skills in problem solving. The Perceptual Reasoning index measures the ability to detect the underlying conceptual relationship among visual objects and use reasoning to identify and apply rules. The Working Memory index measures short-term memory with auditory and visual stimuli. The Processing Speed index measures speed of mental operations and visual-motor coordination. Lastly, the Full-Scale intelligence quotient (IQ) represents a global estimate of intellectual or cognitive ability. All instruments were administered by researchers for the purpose of this research and not professionally or clinically (i.e., for diagnostic or instructional purposes).

### fMRI Tasks

In the scanner, participants performed two different physics tasks, each probing different aspects of physics learning and problem solving (Figure 1).

Participants completed three runs of a *physics reasoning* task, which uses questions from the Force Concept Inventory (FCI; ^79^) to assess domain-specific problem solving. This task includes two conditions, FCI and control, presented in a block design with self-paced trials. FCI and control questions were presented in three screens (Figure 1A, SI Figure 3), between which participants advanced by the press of a button. The first screen presented a written description of a physical scenario and corresponding figure; the second, a question relating to the scenario; and the third, four answer choices from which the participants were instructed choose the correct answer while mentally justifying their choice.

Participants additionally completed two runs of a *physics knowledge* task, which probed physics-related memory retrieval and included physics, general, and control conditions. Participants were asked a series of multiple-choice questions and instructed to respond by indicating their choice with the press of a button (Figure 1B). In the physics condition, participants were asked to recall definitions and formulas taught in the physics course (e.g., “What does the ‘SI’ in SI units stand for?” or “What is the value of the acceleration due to gravity?”). In the general condition, participants were asked to recall general trivia (e.g., “Which of these is not an automobile brand?” or, “Who is the President of the United States?”). The low-level control condition asked participants to press the button corresponding to a letter or symbol. Conditions were organized into blocks and each run included three blocks per condition.

### fMRI Acquisition and Pre-Processing

Neuroimaging data were acquired on a GE 3T Healthcare Discovery 750W MRI scanner at the University of Miami. Functional MRI (fMRI) data were acquired with an interleaved gradient-echo, echo planar imaging (EPI) sequence (TR/TE = 2000/30ms, flip angle = 75°, field of view [FOV] = 220×220mm, matrix size = 64×64, voxel dimensions = 3.4×3.4×3.4mm, 42 axial oblique slices). A T1-weighted series was also acquired using a 3D fast spoiled gradient recall brain volume (FSPGR BRAVO) sequence with 186 contiguous sagittal slices (TI = 650ms, bandwidth = 25.0kHz, flip angle = 12°, FOV = 256×256mm, and slice thickness = 1.0mm). A 2-mm isotropic MNI152 template image was nonlinearly oriented to each participant’s structural T1-weighted image using FMRIB’s Software Library’s (FSL; https://fsl.fmrib.ox.ac.uk/fsl/fslwiki^130^) nonlinear registration tool (FNIRT^131^). Then, each participant’s T1-weighted image was coregistered to the middle volume of each functional run, using FSL’s linear registration tool (FLIRT^132^). These two transformations were concatenated and used to align regionwise parcellations to each subject’s functional images. Tissue-type masks for white matter, gray matter, and cerebrospinal fluid (CSF) were created from each subject’s T1-weighted images using FSL’s automated segmentation tool (FAST^133^).

Task-based fMRI preprocessing began with FSL’s MCFLIRT with spline interpolation, per run per functional task, to align all volumes of each subject’s fMRI time series with that middle volume. To further correct for in-scanner motion effects, functional volumes unduly affected by motion were identified using fsl_motion_outliers, with a framewise displacement threshold of 0.9mm for functional scans.^134^ Data were standardized, detrended, and high-pass filtered, according to the period of each task. The physics knowledge task was high-pass filtered at 0.018Hz and the physics reasoning task was thresholded according to each participant’s individual timing.

### Regionwise Parcellation and Brain Connectivity Analyses

Each participant’s fMRI data were parcellated according to two functionally-derived, whole-brain parcellations with similar numbers of regions. Here, we used a 268-region parcellation computed via multigraph k-way clustering, without spatial constraints, henceforth referred to as the Shen parcellation.^135^ To ensure that our results were not artifacts of node definition, we additionally performed all analyses with an atlas generated from resting-state fMRI data by performing normalized-cut spectral clustering on voxelwise functional connectivity data to define homogeneous, spatially-constrained clusters (i.e., regions), henceforth referred to as the Craddock parcellation,^136^ which includes a range of atlas sizes from which we chose a 268-region solution, to match the granularity of the Shen parcellation. Connectivity graphs were computed using two different brain parcellations (Supplementary Figure 4). Following preprocessing, data analysis continued in two parallel streams, one with the Craddock parcellation and one with the Shen parcellation, to ensure that any results were not artifacts of parcellation schema. Presented results (e.g., Figures 4, 5) include brain-IQ relations observed for both parcellations, but values displayed are derived from the Shen parcellation.

For each region, a single time series was computed as an average of the fMRI time series from all voxels within the region, after further regressing out six motion parameters (from MCFLIRT) and censoring high-motion volumes (framewise displacement >0.9mm), as well as the immediately preceding volume and two following volumes, following recommendations from Power et al. ^137^. Functional tasks’ regionwise time series were standardized (i.e., z-scored), divided by condition per task per run and spliced together across runs, creating separate time series per condition per task for each participant. Adjacency matrices were constructed with each parcellation per participant, per functional task, per session (pre- and post-instruction) using Nilearn (v. 0.3.1, http://nilearn.github.io/index.html), a Python (v 2.7.13) module, built on scikit-learn, for the statistical analysis of neuroimaging data,^138,139^ by computing the pairwise Pearson’s correlations between each pair of regions, resulting in a 268×268 region-wise correlation matrix for each subject per condition per task per session (pre- and post-instruction). Graph theoretic, topological measures were calculated across a range of density-based thresholds. The lowest thresholds at which each network became (a) scale-free and (b) became node-connected were calculated for each adjacency matrix. From these values, a lower- and upper-bound for network thresholding were estimated, following recommendations from Lynall^140^ and Ginestet^141^, such that networks would remain node-connected, meaning there are no brain regions completely separate from the rest of the brain, and spurious connections would be removed while maintaining the scale-free degree distribution expected of the brain per prior research.^142–144^

All topological measures were calculated using bctpy, a Python toolbox intended to replicate the functionality of the Brain Connectivity Toolbox, a MATLAB toolbox for graph theoretic analysis of functional and structural brain connectivity (brain-connectivity-toolbox.net^111^). From each correlation matrix described above, we calculated global efficiency, characteristic path length, and modularity for across the range of proportional thresholds as calculated above (*τ* = [0.21, 0.31] at steps of 0.01), then calculated the area under the curve (AUC) of each measure.^145,146^ These AUCs were used in all following statistical tests assessing the relationship between brain network organization and IQ and will henceforth be referred to per the topological measure from which they were calculated. All topology-related results reported here are significant per topology values calculated from graphs generated from both parcellations.

### Statistical Analyses

Statistical inference was performed using the lavaan R package, and the Python modules SciPy (v. 1.2.1; scipy.org^147–153^) statsmodels (v. 0.9.0; statsmodels.org), and nilearn (v. 0.6.2; nilearn.github.io).

Paired t-tests were used to assess changes in WAIS-IV scores pre-to post-instruction. Two-sample t-tests were used to assess differences in the changes in WAIS-IV scores (post-minus pre-instruction) between male and female students, as well as between students enrolled in the active learning and lecture classes.

Ordinary least squares (OLS) regressions implemented in R were used to regress measures of academic and task performance on WAIS-IV scores, sex, class, age, and years in university, per Equation 1. Significance of individual models was assessed by comparing models’ p-values to a significance threshold adjusted for multiple comparisons via Šidák correction.^81^

A similar procedure, using mass-univariate OLS regressions with permutation testing as implemented by the Python package Nilearn,^154–156^ was used to regress topological measures and functional connectivity (thresholded at *τ* = 0.31) on WAIS-IV scores, sex, class, age, years in university, head size, average framewise displacement (calculated by fsl_motion_outliers, per run, per task), per Equation 2. To correct for multiple comparisons across these regressions, we used the Šidák correction as mentioned above. All reported results were significant in both parcellations, to minimize the effects of brain parcellation on our interpretations.

Mediation models to assess whether brain connectivity explained the relationship between WAIS-IV scores and task performance were run using the R package lavaan.^157^

## Data Availability

A GitHub repository was created at github.com/62442katieb/physics-learning-iq to archive the code and source files for this study, including data preprocessing and analysis scripts and behavioral data. Significant neuroimaging results are available at neurovault.org/collections/9385/.

## Acknowledgments

Primary funding for this project was provided by NSF REAL DRL-1420627; additional support was provided by NSF 1631325, NIH R01-DA041353, NIH U01-DA041156, NSF CNS 1532061, NIH K01-DA037819, NIH U54-MD012393, and the FIU Graduate School Dissertation Year Fellowships (KLB, JEB, RO, AN). We would like to thank the FIU Instructional & Research Computing Center (IRCC, http://ircc.fiu.edu) for providing the HPC and computing resources that contributed to the research results reported within this paper, and to the Department of Psychology of the University of Miami for providing access to their MRI scanner. Special thanks to Karina Falcone, Rosario Pintos Lobo, Emily Boeving, and Camila Uzcategui for their assistance with data collection and to the FIU undergraduate students who volunteered and participated in this project.

## Author Contributions

ARL, EB, SMP, MTS, RWL conceived and designed the project. JEB, EIB, RO, AN acquired behavioral and fMRI data. KLB, JEB, ARL analyzed data. MCR, TS contributed scripts and pipelines. KLB, JEB, ARL wrote the paper and all authors contributed to the revisions and approved the final version.

