## Supplemental Information for "Intelligence and academic performance: Is it all in your head?"

This document includes:

- Supplemental Tables 1 to 7
- Supplemental Figures 1 to 4

#### Supplemental Tables

Supplementary Table 1. F-statistics from ANOVAs assessing pre- to post-instruction changes in WAIS-IV scores with respect to students' sex and classroom environment

| WAIS-IV Score | Time (post>pre) | Sex X Time | Class X Time | Sex X Class X Time |
|---|---|---|---|---|
| FSIQ | 71.390** | 2.001 | 3.577 | 0.066 |
| PSI | 47.856** | 0.180 | 7.327* | 0.722 |
| PRI | 30.856** | 1.162 | 0.405 | 0.123 |
| VCI | 4.925 | 0.616 | 0.063 | 0.353 |
| WMI | 2.974 | 1.083 | 0.545 | 0.506 |

*Indicates statistic significant at $p_{FWE\text{-}corr} < 0.05$; ** indicates statistic significant at $p_{FWE\text{-}corr} < 0.01$.

Supplementary Table 2. Results of all OLS regressions assessing relationships between performance and WAIS-IV scores, without WAIS-IV interactions.

| | F | p(F) | BIC | IQ | p(IQ) | Sex | p(Sex) | Class | p(Class) | Age | p(Age) | Year in Univ. | p(Year in Univ) |
|---|---|---|---|---|---|---|---|---|---|---|---|---|---|
| Course grade | | | | | | | | | | | | | |
| deltaFSIQ | 1.52 | 1.78E-01 | 387 | -0.004 | 7.32E-01 | 0.306 | 2.12E-01 | 0.379 | 1.22E-01 | -0.128 | 1.44E-01 | 0.129 | 3.02E-01 |
| deltaPRI | 2.24 | 4.39E-02 | 379 | 0.015 | 6.50E-02 | 0.300 | 2.18E-01 | 0.411 | 8.24E-02 | -0.098 | 2.38E-01 | 0.113 | 3.60E-01 |
| deltaPSI | 1.65 | 1.38E-01 | 386 | -0.005 | 4.05E-01 | 0.310 | 2.09E-01 | 0.419 | 1.07E-01 | -0.123 | 1.47E-01 | 0.123 | 3.21E-01 |
| deltaVCI | 1.60 | 1.53E-01 | 392 | -0.007 | 4.96E-01 | 0.306 | 2.11E-01 | 0.381 | 1.23E-01 | -0.126 | 1.42E-01 | 0.127 | 3.02E-01 |
| deltaWMI | 2.05 | 6.42E-02 | 387 | -0.016 | 7.53E-02 | 0.317 | 1.93E-01 | 0.329 | 1.71E-01 | -0.146 | 7.03E-02 | 0.150 | 2.35E-01 |
| FSIQ2 | 1.55 | 1.68E-01 | 389 | 0.005 | 5.83E-01 | 0.278 | 2.60E-01 | 0.378 | 1.23E-01 | -0.113 | 1.99E-01 | 0.117 | 3.53E-01 |
| PRI2 | 1.87 | 9.20E-02 | 386 | 0.012 | 1.78E-01 | 0.242 | 3.10E-01 | 0.388 | 1.08E-01 | -0.110 | 1.76E-01 | 0.129 | 2.88E-01 |
| PSI2 | 1.67 | 1.35E-01 | 381 | -0.006 | 3.64E-01 | 0.294 | 2.24E-01 | 0.416 | 1.00E-01 | -0.129 | 1.37E-01 | 0.127 | 2.97E-01 |
| VCI2 | 1.61 | 1.51E-01 | 391 | 0.007 | 5.23E-01 | 0.266 | 3.10E-01 | 0.407 | 1.04E-01 | -0.109 | 2.05E-01 | 0.108 | 3.86E-01 |
| WMI2 | 1.49 | 1.86E-01 | 390 | 0.000 | 9.83E-01 | 0.305 | 2.23E-01 | 0.374 | 1.31E-01 | -0.123 | 1.56E-01 | 0.127 | 3.11E-01 |
| Physics knowledge accuracy | | | | | | | | | | | | | |
| deltaFSIQ | 3.81 | **1.62E-03** | -173 | 0.000 | 9.49E-01 | 0.063 | 3.25E-02 | -0.047 | 8.39E-02 | -0.009 | 2.75E-01 | -0.004 | 7.64E-01 |

| | | | | | | | | | | | | | |
|---|---|---|---|---|---|---|---|---|---|---|---|---|---|
| deltaPRI | 4.08 | **9.12E-04** | -175 | 0.001 | 2.31E-01 | 0.063 | 3.25E-02 | -0.044 | 1.00E-01 | -0.008 | 3.41E-01 | -0.005 | 7.13E-01 |
| deltaPSI | 3.81 | **1.62E-03** | -174 | 0.000 | 9.09E-01 | 0.063 | 3.28E-02 | -0.047 | 1.04E-01 | -0.010 | 2.46E-01 | -0.004 | 7.74E-01 |
| deltaVCI | 4.42 | **4.47E-04** | -174 | -0.002 | 1.43E-01 | 0.063 | 2.99E-02 | -0.045 | 9.24E-02 | -0.010 | 2.32E-01 | -0.004 | 7.77E-01 |
| deltaWMI | 3.82 | **1.61E-03** | -173 | 0.000 | 8.63E-01 | 0.063 | 3.26E-02 | -0.046 | 9.33E-02 | -0.009 | 2.74E-01 | -0.004 | 7.57E-01 |
| **FSIQ2** | 5.85 | **2.16E-05** | -185 | 0.003 | **4.65E-03** | 0.047 | 1.23E-01 | -0.044 | 8.98E-02 | -0.003 | 7.01E-01 | -0.009 | 4.44E-01 |
| PRI2 | 3.81 | **1.62E-03** | -177 | 0.000 | 9.19E-01 | 0.062 | 3.15E-02 | -0.046 | 8.67E-02 | -0.009 | 2.54E-01 | -0.004 | 7.70E-01 |
| PSI2 | 3.83 | **1.54E-03** | -176 | 0.000 | 7.50E-01 | 0.063 | 3.22E-02 | -0.048 | 9.76E-02 | -0.009 | 2.59E-01 | -0.004 | 7.67E-01 |
| **VCI2** | 7.83 | **3.86E-07** | -197 | 0.004 | **1.46E-03** | 0.040 | 1.80E-01 | -0.027 | 3.19E-01 | -0.001 | 8.88E-01 | -0.015 | 2.30E-01 |
| **WMI2** | 6.39 | **7.10E-06** | -188 | 0.003 | **4.40E-04** | 0.047 | 1.13E-01 | -0.044 | 1.00E-01 | -0.004 | 6.52E-01 | -0.013 | 2.86E-01 |
| Physics reasoning accuracy | | | | | | | | | | | | | |
| deltaFSIQ | 5.26 | **7.46E-05** | -60 | 0.003 | 1.92E-01 | 0.158 | 1.37E-04 | -0.022 | 6.06E-01 | -0.023 | 4.92E-02 | 0.011 | 6.42E-01 |
| **deltaPRI** | 5.83 | **2.29E-05** | -63 | 0.003 | **3.93E-02** | 0.159 | 1.20E-04 | -0.012 | 7.77E-01 | -0.021 | 6.15E-02 | 0.010 | 6.70E-01 |
| deltaPSI | 4.94 | **1.48E-04** | -51 | 0.000 | 8.72E-01 | 0.159 | 1.08E-04 | -0.020 | 6.64E-01 | -0.026 | 2.13E-02 | 0.012 | 5.96E-01 |
| deltaVCI | 4.94 | **1.49E-04** | -51 | 0.000 | 8.82E-01 | 0.160 | 9.44E-05 | -0.019 | 6.67E-01 | -0.026 | 1.96E-02 | 0.012 | 5.97E-01 |
| deltaWMI | 4.96 | **1.42E-04** | -55 | -0.001 | 6.96E-01 | 0.160 | 8.49E-05 | -0.020 | 6.41E-01 | -0.027 | 1.45E-02 | 0.013 | 5.71E-01 |

| | | | | | | | | | | | | | |
|---|---|---|---|---|---|---|---|---|---|---|---|---|---|
| **FSIQ2** | 8.56 | **9.09E-08** | -69 | 0.007 | **1.70E-04** | 0.127 | 2.56E-03 | -0.014 | 7.29E-01 | -0.013 | 2.40E-01 | 0.000 | 9.84E-01 |
| **PRI2** | 6.75 | **3.38E-06** | -64 | 0.004 | **4.33E-04** | 0.137 | 8.20E-04 | -0.014 | 7.29E-01 | -0.021 | 3.49E-02 | 0.013 | 5.55E-01 |
| PSI2 | 4.96 | **1.40E-04** | -54 | 0.000 | 7.28E-01 | 0.160 | 9.93E-05 | -0.022 | 6.29E-01 | -0.025 | 2.59E-02 | 0.012 | 6.06E-01 |
| **VCI2** | 10.58 | **1.96E-09** | -76 | 0.007 | **5.58E-05** | 0.119 | 4.05E-03 | 0.016 | 6.90E-01 | -0.011 | 3.01E-01 | -0.008 | 7.33E-01 |
| WMI2 | 5.36 | **6.12E-05** | -54 | 0.002 | 1.88E-01 | 0.149 | 3.26E-04 | -0.017 | 6.91E-01 | -0.022 | 4.38E-02 | 0.006 | 7.84E-01 |

Bold WAIS-IV scores indicate a model in which a WAIS-IV score and/or WAIS-IV interaction term is significantly related to the performance measure, i.e., that the model is significant at $p_{FWE\text{-}corr} < 0.05$ and the WAIS-IV score regression coefficient is significant at $p < 0.05$. Bold model F-statistics indicate that the model is significant at $p_{FWE\text{-}corr} < 0.05$. Bold p-values indicate significant regression coefficients of significant relationships at $p < 0.05$. All model fit statistics (i.e., P(F)) were assessed for significance at a FWE-corrected $\alpha < 0.05$.

**Supplementary Table 3.** Results of all OLS regressions assessing relationships between performance and WAIS-IV scores, with WAIS-IV interactions.

| | F | p(F) | BIC | IQ | p(IQ) | IQ *Sex | p(IQ *Sex) | IQ* Class | p(IQ* Class) | IQ *Sex*Class | p(IQ *Sex* Class) | Sex | p(Sex) | Class | p(Class) | Sex* Class | p(Sex* Class) | Age | p(Age) | Year in Univ. | p(Year in Univ.) | ANOVA vs. Simple |
|---|---|---|---|---|---|---|---|---|---|---|---|---|---|---|---|---|---|---|---|---|---|---|
| Course grade | | | | | | | | | | | | | | | | | | | | | | |
| deltaFSIQ | 1.59 | 1.26E-01 | 387 | 0.01 | 5.33E-01 | -0.05 | 1.12E-01 | -0.03 | 3.72E-01 | 0.09 | 5.38E-02 | 0.60 | **4.16E-02** | 0.58 | 7.29E-02 | -0.77 | 1.15E-01 | -0.14 | 1.29E-01 | 0.14 | 2.56E-01 | 0.17 |
| deltaPRI | 2.56 | 9.99E-03 | 379 | 0.04 | **1.63E-02** | -0.06 | **6.64E-03** | -0.01 | 6.25E-01 | 0.04 | 1.46E-01 | 0.73 | **2.38E-02** | 0.54 | 6.83E-02 | -0.56 | 1.91E-01 | -0.12 | 1.49E-01 | 0.16 | 2.19E-01 | **0.03** |
| deltaPSI | 1.74 | 8.66E-02 | 386 | -0.01 | 5.00E-01 | -0.01 | 5.54E-01 | 0.00 | 9.78E-01 | 0.03 | 1.45E-01 | 0.41 | 1.41E-01 | 0.47 | 1.59E-01 | -0.59 | 1.93E-01 | -0.13 | 1.31E-01 | 0.11 | 3.72E-01 | 0.14 |
| deltaVCI | 1.10 | 3.66E-01 | 392 | -0.01 | 7.89E-01 | -0.01 | 8.15E-01 | 0.01 | 7.17E-01 | -0.01 | 8.23E-01 | 0.32 | 1.98E-01 | 0.34 | 2.13E-01 | -0.10 | 7.90E-01 | -0.13 | 1.38E-01 | 0.13 | 3.06E-01 | 0.92 |
| deltaWMI | 1.66 | 1.07E-01 | 387 | 0.00 | 8.39E-01 | 0.00 | 9.86E-01 | -0.04 | 1.53E-01 | 0.02 | 5.55E-01 | 0.31 | 2.43E-01 | 0.32 | 1.94E-01 | -0.01 | 9.70E-01 | -0.15 | 7.07E-02 | 0.15 | 2.35E-01 | 0.45 |
| FSIQ2 | 1.43 | 1.82E-01 | 389 | 0.03 | 1.70E-01 | -0.05 | 5.77E-02 | -0.03 | 3.29E-01 | 0.07 | 8.56E-02 | 6.03 | 5.23E-02 | 3.86 | 2.87E-01 | -8.06 | 8.34E-02 | -0.12 | 1.72E-01 | 0.12 | 3.46E-01 | 0.32 |
| PRI2 | 1.69 | 9.81E-02 | 386 | 0.02 | 9.87E-02 | -0.05 | 6.28E-02 | -0.01 | 6.12E-01 | 0.06 | 1.61E-01 | 6.37 | 5.02E-02 | 1.58 | 5.07E-01 | -6.56 | 1.46E-01 | -0.11 | 1.71E-01 | 0.12 | 3.35E-01 | 0.27 |
| PSI2 | 2.35 | 1.76E-02 | 381 | 0.00 | 8.08E-01 | -0.03 | 2.28E-01 | 0.00 | 9.89E-01 | 0.05 | 6.96E-02 | 3.04 | 1.83E-01 | 0.37 | 8.80E-01 | -5.33 | 6.44E-02 | -0.14 | 1.01E-01 | 0.11 | 3.43E-01 | **0.02** |
| VCI2 | 1.16 | 3.30E-01 | 391 | 0.01 | 5.38E-01 | 0.00 | 9.92E-01 | -0.02 | 4.78E-01 | 0.01 | 7.82E-01 | 0.20 | 9.60E-01 | 2.77 | 4.21E-01 | -1.33 | 7.87E-01 | -0.10 | 2.34E-01 | 0.11 | 4.02E-01 | 0.82 |
| WMI2 | 1.26 | 2.64E-01 | 390 | 0.00 | 9.20E-01 | 0.02 | 6.33E-01 | -0.02 | 6.80E-01 | -0.01 | 7.82E-01 | -1.62 | 6.84E-01 | 2.04 | 6.21E-01 | 1.34 | 7.84E-01 | -0.11 | 2.26E-01 | 0.12 | 3.21E-01 | 0.49 |

| | | | | | | | | | | | | | | | | | | | | | | |
|---|---|---|---|---|---|---|---|---|---|---|---|---|---|---|---|---|---|---|---|---|---|---|
| Physics knowledge accuracy | | | | | | | | | | | | | | | | | | | | | | |
| deltaFSIQ | 2.81 | **4.99E-03** | -173 | 0.00 | 2.48E-01 | 0.00 | 3.19E-01 | 0.01 | 9.90E-02 | -0.01 | 1.65E-01 | 0.04 | 3.44E-01 | -0.08 | 3.16E-02 | 0.07 | 2.05E-01 | -0.01 | 2.51E-01 | 0.00 | 7.48E-01 | 0.47 |
| deltaPRI | 3.02 | **2.79E-03** | -175 | 0.00 | 6.49E-01 | 0.00 | 7.75E-01 | 0.00 | 3.64E-01 | -0.01 | 2.24E-01 | 0.06 | 1.67E-01 | -0.06 | 1.22E-01 | 0.05 | 3.04E-01 | -0.01 | 4.09E-01 | -0.01 | 6.25E-01 | 0.44 |
| deltaPSI | 2.98 | **3.15E-03** | -174 | 0.00 | 1.45E-01 | 0.00 | 3.82E-01 | 0.00 | 7.74E-02 | 0.00 | 3.56E-01 | 0.05 | 1.10E-01 | -0.08 | 2.96E-02 | 0.05 | 3.43E-01 | -0.01 | 2.04E-01 | 0.00 | 7.83E-01 | 0.29 |
| deltaVCI | 2.89 | **3.99E-03** | -174 | 0.00 | 5.19E-01 | 0.00 | 9.59E-01 | 0.00 | 8.39E-01 | 0.00 | 9.60E-01 | 0.06 | 3.91E-02 | -0.04 | 1.10E-01 | 0.03 | 4.92E-01 | -0.01 | 2.44E-01 | 0.00 | 8.13E-01 | 0.99 |
| deltaWMI | 2.80 | **5.13E-03** | -173 | 0.00 | 6.18E-01 | 0.00 | 1.54E-01 | 0.00 | 7.93E-01 | 0.00 | 5.76E-01 | 0.05 | 9.39E-02 | -0.05 | 7.22E-02 | 0.04 | 3.23E-01 | -0.01 | 3.84E-01 | -0.01 | 5.81E-01 | 0.49 |
| FSIQ2 | 4.32 | **6.98E-05** | -185 | 0.00 | 8.71E-01 | 0.01 | 6.59E-02 | 0.00 | 3.52E-01 | -0.01 | 1.96E-01 | -0.59 | 1.06E-01 | -0.53 | 3.17E-01 | 0.82 | 1.92E-01 | 0.00 | 6.96E-01 | -0.01 | 4.63E-01 | 0.31 |
| PRI2 | 3.38 | **1.00E-03** | -177 | 0.00 | 4.84E-01 | 0.00 | 2.58E-01 | 0.00 | 3.38E-01 | 0.00 | 7.70E-01 | 0.55 | 1.87E-01 | -0.49 | 3.04E-01 | -0.19 | 7.65E-01 | -0.01 | 2.30E-01 | 0.00 | 7.03E-01 | 0.08 |
| PSI2 | 3.19 | **1.74E-03** | -176 | 0.00 | 1.71E-01 | 0.00 | 5.73E-01 | 0.00 | 7.14E-02 | 0.00 | 3.43E-01 | -0.06 | 7.74E-01 | -0.52 | 5.77E-02 | 0.33 | 3.02E-01 | -0.01 | 1.87E-01 | 0.00 | 8.16E-01 | 0.16 |
| **VCI2** | 6.03 | **6.14E-07** | -197 | 0.00 | **4.18E-02** | 0.00 | 1.53E-01 | 0.00 | 5.33E-01 | 0.00 | 6.45E-01 | -0.44 | 2.07E-01 | 0.14 | 6.27E-01 | 0.27 | 5.80E-01 | 0.00 | 9.34E-01 | -0.01 | 2.52E-01 | 0.11 |
| **WMI2** | 4.74 | **2.12E-05** | -188 | 0.00 | 7.37E-01 | 0.01 | **2.31E-02** | 0.00 | 5.79E-01 | 0.00 | 2.48E-01 | -0.51 | 4.16E-02 | -0.23 | 4.97E-01 | 0.50 | 2.23E-01 | 0.00 | 6.11E-01 | -0.01 | 2.70E-01 | 0.26 |
| Physics reasoning accuracy | | | | | | | | | | | | | | | | | | | | | | |
| **deltaFSIQ** | 4.50 | **4.24E-05** | -60 | 0.00 | **3.77E-02** | -0.01 | **2.81E-03** | 0.00 | 8.87E-01 | 0.01 | 1.56E-01 | 0.22 | **1.07E-06** | -0.03 | 6.41E-01 | -0.12 | 1.78E-01 | -0.03 | **1.98E-02** | 0.02 | 4.89E-01 | 0.06 |
| **deltaPRI** | 5.03 | **9.42E-06** | -63 | 0.01 | **7.11E-03** | -0.01 | **9.58E-04** | 0.00 | 8.87E-01 | 0.01 | 2.51E-01 | 0.23 | **1.49E-06** | 0.00 | 9.15E-01 | -0.11 | 1.22E-01 | -0.03 | **1.50E-02** | 0.02 | 3.53E-01 | **0.04** |
| deltaPSI | 3.40 | **9.38E-04** | -51 | 0.00 | 6.85E-01 | 0.00 | 9.38E-01 | 0.00 | 7.16E-01 | 0.00 | 8.02E-01 | 0.16 | **7.68E-04** | -0.03 | 6.71E-01 | -0.04 | 6.51E-01 | -0.03 | **2.55E-02** | 0.01 | 6.24E-01 | 0.71 |

| | | | | | | | | | | | | | | | | | | | | | | |
|---|---|---|---|---|---|---|---|---|---|---|---|---|---|---|---|---|---|---|---|---|---|---|
| deltaVCI | 3.35 | **1.10E-03** | -51 | 0.00 | 5.29E-01 | 0.00 | 3.08E-01 | 0.00 | 4.98E-01 | 0.01 | 3.16E-01 | 0.17 | **7.76E-05** | -0.01 | 8.08E-01 | -0.04 | 5.08E-01 | -0.02 | **2.75E-02** | 0.01 | 6.13E-01 | 0.80 |
| deltaWMI | 3.87 | **2.53E-04** | -55 | 0.00 | 9.81E-01 | 0.00 | 3.29E-01 | 0.01 | 2.42E-01 | 0.00 | 7.44E-01 | 0.17 | **5.39E-05** | -0.01 | 8.12E-01 | -0.04 | 5.14E-01 | -0.03 | **1.23E-02** | 0.02 | 4.37E-01 | 0.21 |
| **FSIQ2** | 5.77 | **1.24E-06** | -69 | 0.01 | **4.95E-02** | 0.00 | 5.89E-01 | 0.00 | 5.88E-01 | 0.00 | 9.14E-01 | 0.46 | 4.54E-01 | -0.32 | 5.76E-01 | 0.04 | 9.63E-01 | -0.01 | 1.78E-01 | 0.00 | 9.62E-01 | 0.73 |
| **PRI2** | 5.12 | **7.40E-06** | -64 | 0.01 | **1.50E-03** | -0.01 | **2.48E-02** | 0.00 | 5.54E-01 | 0.01 | 2.60E-01 | 1.19 | **1.16E-02** | -0.21 | 5.29E-01 | -0.76 | 2.28E-01 | -0.02 | **2.58E-02** | 0.01 | 6.29E-01 | 0.18 |
| PSI2 | 3.79 | **3.14E-04** | -54 | 0.00 | 4.65E-01 | 0.00 | 7.40E-01 | 0.01 | 2.45E-01 | 0.00 | 7.30E-01 | 0.01 | 9.89E-01 | -0.62 | 2.34E-01 | 0.19 | 7.58E-01 | -0.03 | **1.86E-02** | 0.01 | 6.05E-01 | 0.26 |
| VCI2 | 6.89 | **6.13E-08** | -76 | 0.01 | 1.59E-01 | 0.00 | 9.00E-01 | 0.00 | 9.34E-01 | 0.00 | 9.65E-01 | 0.05 | 9.35E-01 | -0.05 | 9.51E-01 | 0.00 | 9.99E-01 | -0.01 | 2.89E-01 | -0.01 | 7.48E-01 | 1.00 |
| WMI2 | 3.71 | **3.90E-04** | -54 | 0.00 | 7.98E-01 | 0.00 | 8.68E-01 | 0.00 | 4.85E-01 | -0.01 | 3.97E-01 | 0.04 | 9.43E-01 | -0.44 | 4.80E-01 | 0.61 | 4.24E-01 | -0.02 | 6.05E-02 | 0.01 | 7.56E-01 | 0.65 |

icantly related to the performance measure, i.e., that the model is significant at $p_{FWE\text{-}corr} < 0.05$ and the WAIS-IV score regression coefficient is significant at $p < 0.05$. Bold model F-statistics indicate that the model is significant at $p_{FWE\text{-}corr} < 0.05$. Bold p-values indicate significant regression coefficients of significant relationships at $p < 0.05$. All model fit statistics (i.e., P(F)) were assessed for significance at a FWE-corrected $\alpha < 0.05$. Italicized WAIS-IV scores indicate models that explain significantly more variance than the nested model without WAIS-interaction terms at $p < 0.05$.

**Supplementary Table 4.** Results of all OLS regressions assessing relationships between brain network efficiency and WAIS-IV scores, without WAIS-IV interactions.

| WAIS-IV Score | F | p(F) | BIC | WAIS | p(WAIS) | Sex | p(Sex) | Class | p(Class) | Age | p(Age) | Year in Univ. | p(Year in Univ.) |
|---|---|---|---|---|---|---|---|---|---|---|---|---|---|
| Physics knowledge | | | | | | | | | | | | | |
| Craddock et al., (2012) | | | | | | | | | | | | | |
| deltaFSIQ | 0.389 | 8.85E-01 | -2 | 0.002 | 3.78E-01 | 0.019 | 7.84E-01 | -0.015 | 8.09E-01 | 0.002 | 9.31E-01 | 0.028 | 4.36E-01 |
| deltaPRI | 0.337 | 9.16E-01 | -10 | 0.001 | 5.79E-01 | 0.024 | 7.38E-01 | -0.007 | 9.16E-01 | 0.000 | 9.99E-01 | 0.029 | 4.25E-01 |
| deltaPSI | 0.278 | 9.46E-01 | -1 | 0.000 | 7.42E-01 | 0.029 | 6.84E-01 | -0.003 | 9.70E-01 | -0.003 | 8.76E-01 | 0.030 | 4.17E-01 |
| deltaVCI | 0.463 | 8.34E-01 | -1 | 0.002 | 3.32E-01 | 0.022 | 7.64E-01 | -0.013 | 8.33E-01 | -0.001 | 9.56E-01 | 0.030 | 4.20E-01 |
| deltaWMI | 0.381 | 8.90E-01 | 3 | 0.002 | 4.12E-01 | 0.023 | 7.43E-01 | -0.003 | 9.59E-01 | 0.000 | 9.87E-01 | 0.027 | 4.51E-01 |
| FSIQ2 | 0.294 | 9.38E-01 | -7 | 0.001 | 6.93E-01 | 0.020 | 7.76E-01 | -0.007 | 9.13E-01 | 0.000 | 9.92E-01 | 0.028 | 4.40E-01 |
| PRI2 | 0.342 | 9.13E-01 | -10 | 0.001 | 5.92E-01 | 0.017 | 8.11E-01 | -0.008 | 9.06E-01 | -0.001 | 9.62E-01 | 0.030 | 4.10E-01 |
| PSI2 | 0.256 | 9.56E-01 | 3 | 0.000 | 9.47E-01 | 0.027 | 7.06E-01 | -0.009 | 9.01E-01 | -0.002 | 9.02E-01 | 0.030 | 4.18E-01 |
| VCI2 | 0.259 | 9.55E-01 | 3 | 0.000 | 8.87E-01 | 0.025 | 7.32E-01 | -0.006 | 9.28E-01 | -0.002 | 9.29E-01 | 0.029 | 4.55E-01 |
| WMI2 | 0.256 | 9.56E-01 | -1 | 0.000 | 9.70E-01 | 0.027 | 7.04E-01 | -0.008 | 9.01E-01 | -0.002 | 8.96E-01 | 0.030 | 4.21E-01 |
| Shen et al., (2015) | | | | | | | | | | | | | |

| | | | | | | | | | | | | | |
|---|---|---|---|---|---|---|---|---|---|---|---|---|---|
| deltaFSIQ | 0.405 | 8.74E-01 | 15 | 0.003 | 3.29E-01 | 0.023 | 7.55E-01 | -0.018 | 7.90E-01 | 0.005 | 8.28E-01 | 0.026 | 5.12E-01 |
| deltaPRI | 0.326 | 9.22E-01 | 7 | 0.001 | 5.75E-01 | 0.030 | 7.04E-01 | -0.008 | 9.07E-01 | 0.002 | 9.11E-01 | 0.027 | 4.98E-01 |
| deltaPSI | 0.259 | 9.54E-01 | 15 | 0.000 | 7.79E-01 | 0.035 | 6.53E-01 | -0.004 | 9.53E-01 | -0.001 | 9.74E-01 | 0.028 | 4.89E-01 |
| deltaVCI | 0.481 | 8.21E-01 | 16 | 0.003 | 2.89E-01 | 0.027 | 7.32E-01 | -0.015 | 8.17E-01 | 0.001 | 9.48E-01 | 0.028 | 4.91E-01 |
| deltaWMI | 0.390 | 8.84E-01 | 20 | 0.002 | 3.78E-01 | 0.029 | 7.11E-01 | -0.004 | 9.54E-01 | 0.003 | 8.89E-01 | 0.025 | 5.30E-01 |
| FSIQ2 | 0.255 | 9.56E-01 | 11 | 0.001 | 8.29E-01 | 0.029 | 7.13E-01 | -0.009 | 9.00E-01 | 0.001 | 9.59E-01 | 0.027 | 5.03E-01 |
| PRI2 | 0.291 | 9.40E-01 | 8 | 0.001 | 6.97E-01 | 0.025 | 7.51E-01 | -0.009 | 8.97E-01 | 0.001 | 9.62E-01 | 0.028 | 4.84E-01 |
| PSI2 | 0.243 | 9.61E-01 | 19 | 0.000 | 9.80E-01 | 0.033 | 6.71E-01 | -0.009 | 9.00E-01 | 0.000 | 9.95E-01 | 0.028 | 4.91E-01 |
| VCI2 | 0.243 | 9.61E-01 | 20 | 0.000 | 9.81E-01 | 0.033 | 6.87E-01 | -0.009 | 9.00E-01 | 0.000 | 9.97E-01 | 0.028 | 5.15E-01 |
| WMI2 | 0.254 | 9.56E-01 | 16 | -0.001 | 7.86E-01 | 0.036 | 6.53E-01 | -0.010 | 8.80E-01 | -0.001 | 9.51E-01 | 0.030 | 4.71E-01 |
| Physics reasoning | | | | | | | | | | | | | |
| Craddock et al., (2012) | | | | | | | | | | | | | |
| deltaFSIQ | 0.263 | 9.53E-01 | -20 | 0.001 | 8.46E-01 | 0.035 | 5.79E-01 | -0.014 | 8.06E-01 | 0.012 | 5.06E-01 | -0.010 | 7.80E-01 |
| deltaPRI | 0.405 | 8.74E-01 | -29 | 0.002 | 4.21E-01 | 0.031 | 6.17E-01 | -0.012 | 8.33E-01 | 0.014 | 4.31E-01 | -0.010 | 7.80E-01 |
| deltaPSI | 0.292 | 9.39E-01 | -16 | 0.000 | 6.76E-01 | 0.036 | 5.85E-01 | -0.018 | 7.60E-01 | 0.011 | 5.29E-01 | -0.010 | 7.79E-01 |

| | | | | | | | | | | | | | |
|---|---|---|---|---|---|---|---|---|---|---|---|---|---|
| deltaVCI | 0.273 | 9.48E-01 | -19 | -0.001 | 7.77E-01 | 0.038 | 5.64E-01 | -0.011 | 8.46E-01 | 0.010 | 5.70E-01 | -0.010 | 7.89E-01 |
| deltaWMI | 0.388 | 8.85E-01 | -16 | -0.002 | 4.69E-01 | 0.041 | 5.30E-01 | -0.016 | 7.81E-01 | 0.008 | 6.44E-01 | -0.008 | 8.38E-01 |
| FSIQ2 | 0.291 | 9.40E-01 | -25 | -0.001 | 6.81E-01 | 0.043 | 5.29E-01 | -0.013 | 8.23E-01 | 0.009 | 6.23E-01 | -0.009 | 8.19E-01 |
| PRI2 | 0.265 | 9.52E-01 | -28 | 0.000 | 8.42E-01 | 0.034 | 6.03E-01 | -0.013 | 8.26E-01 | 0.011 | 5.36E-01 | -0.010 | 7.97E-01 |
| PSI2 | 0.262 | 9.53E-01 | -16 | 0.000 | 8.62E-01 | 0.037 | 5.75E-01 | -0.015 | 8.09E-01 | 0.011 | 5.42E-01 | -0.010 | 7.91E-01 |
| VCI2 | 0.409 | 8.71E-01 | -17 | -0.002 | 3.49E-01 | 0.048 | 5.02E-01 | -0.023 | 6.92E-01 | 0.007 | 7.01E-01 | -0.005 | 8.91E-01 |
| WMI2 | 0.379 | 8.91E-01 | -25 | -0.002 | 4.33E-01 | 0.044 | 5.15E-01 | -0.015 | 7.90E-01 | 0.008 | 6.65E-01 | -0.006 | 8.83E-01 |
| Shen et al., (2015) | | | | | | | | | | | | | |
| deltaFSIQ | 0.208 | 9.73E-01 | -7 | 0.001 | 8.31E-01 | 0.030 | 6.55E-01 | -0.017 | 7.78E-01 | 0.010 | 5.86E-01 | -0.011 | 7.76E-01 |
| deltaPRI | 0.307 | 9.32E-01 | -16 | 0.001 | 5.00E-01 | 0.027 | 6.86E-01 | -0.015 | 8.08E-01 | 0.012 | 5.26E-01 | -0.011 | 7.80E-01 |
| deltaPSI | 0.217 | 9.71E-01 | -3 | 0.000 | 7.70E-01 | 0.031 | 6.56E-01 | -0.020 | 7.60E-01 | 0.010 | 6.22E-01 | -0.011 | 7.80E-01 |
| deltaVCI | 0.200 | 9.76E-01 | -6 | 0.000 | 9.93E-01 | 0.032 | 6.51E-01 | -0.016 | 8.00E-01 | 0.009 | 6.44E-01 | -0.011 | 7.88E-01 |
| deltaWMI | 0.281 | 9.45E-01 | -3 | -0.001 | 5.57E-01 | 0.036 | 6.11E-01 | -0.018 | 7.65E-01 | 0.007 | 7.12E-01 | -0.009 | 8.24E-01 |
| FSIQ2 | 0.292 | 9.39E-01 | -14 | -0.002 | 5.12E-01 | 0.042 | 5.63E-01 | -0.016 | 7.97E-01 | 0.006 | 7.56E-01 | -0.009 | 8.34E-01 |
| PRI2 | 0.200 | 9.76E-01 | -17 | 0.000 | 9.63E-01 | 0.033 | 6.36E-01 | -0.015 | 8.05E-01 | 0.009 | 6.42E-01 | -0.011 | 7.88E-01 |

| | | | | | | | | | | | | | |
|---|---|---|---|---|---|---|---|---|---|---|---|---|---|
| PSI2 | 0.202 | 9.76E-01 | -3 | 0.000 | 9.26E-01 | 0.032 | 6.48E-01 | -0.017 | 8.00E-01 | 0.009 | 6.32E-01 | -0.011 | 7.88E-01 |
| VCI2 | 0.414 | 8.68E-01 | -6 | -0.002 | 2.68E-01 | 0.046 | 5.47E-01 | -0.028 | 6.46E-01 | 0.005 | 8.12E-01 | -0.005 | 9.02E-01 |
| WMI2 | 0.355 | 9.06E-01 | -13 | -0.002 | 3.58E-01 | 0.041 | 5.75E-01 | -0.019 | 7.60E-01 | 0.006 | 7.73E-01 | -0.006 | 8.90E-01 |

Bold WAIS-IV scores indicate a model in which a WAIS-IV score and/or WAIS-IV interaction term is significantly related to the performance measure, i.e., that the model is significant at $p_{FWE\text{-}corr} < 0.05$ and the WAIS-IV score regression coefficient is significant at $p < 0.05$. Bold model F-statistics indicate that the model is significant at $p_{FWE\text{-}corr} < 0.05$. Bold p-values indicate significant regression coefficients of significant relationships at $p < 0.05$. All model fit statistics (i.e., P(F)) were assessed for significance at a FWE-corrected $\alpha < 0.05$.

**Supplementary Table 5.** Results of all OLS regressions assessing relationships between brain network efficiency and WAIS-IV scores, with WAIS-IV interactions.

| WAIS-IV Score | F | p(F) | BIC | WAIS | p(WAIS) | IQ *Sex | p(IQ *Sex) | IQ* Class | p(IQ* Class) | IQ *Sex*Class | p(IQ *Sex* Class) | Sex | p(Sex) | Class | p(Class) | Sex* Class | p(Sex* Class) | Age | p(Age) | Year in Univ. | p(Year in Univ.) | ANOVA vs. Simple |
|---|---|---|---|---|---|---|---|---|---|---|---|---|---|---|---|---|---|---|---|---|---|---|
| Physics reasoning | | | | | | | | | | | | | | | | | | | | | | |
| Craddock et al., (2012) | | | | | | | | | | | | | | | | | | | | | | |
| deltaFSIQ | 0.86 | 5.6E-01 | -20 | 0.01 | **4.7E-03** | -0.01 | 1.8E-01 | -0.02 | **1.8E-03** | 0.02 | 6.6E-02 | 0.07 | 3.0E-01 | 0.08 | 2.6E-01 | -0.10 | 2.9E-01 | 0.02 | 3.1E-01 | -0.01 | 8.3E-01 | 0.11 |
| deltaPRI | 1.87 | 6.7E-02 | -29 | 0.00 | 1.1E-01 | 0.00 | 7.5E-01 | -0.02 | **8.7E-04** | 0.01 | 1.2E-01 | 0.00 | 9.7E-01 | 0.06 | 3.4E-01 | -0.04 | 6.0E-01 | 0.02 | 2.2E-01 | -0.02 | 5.9E-01 | **0.00** |
| deltaPSI | 0.45 | 9.1E-01 | -16 | 0.00 | 2.2E-01 | 0.00 | 6.3E-01 | 0.00 | 4.5E-01 | 0.00 | 9.8E-01 | 0.04 | 5.0E-01 | 0.00 | 9.6E-01 | 0.00 | 9.8E-01 | 0.01 | 4.9E-01 | -0.01 | 8.1E-01 | 0.52 |
| deltaVCI | 0.72 | 6.9E-01 | -19 | 0.00 | 8.1E-01 | -0.01 | 1.1E-01 | 0.00 | 8.4E-01 | 0.01 | 3.8E-01 | 0.05 | 4.1E-01 | -0.02 | 7.6E-01 | -0.03 | 7.0E-01 | 0.01 | 5.8E-01 | -0.02 | 6.9E-01 | 0.19 |
| deltaWMI | 0.45 | 9.0E-01 | -16 | 0.00 | 5.3E-01 | 0.00 | 6.0E-01 | -0.01 | 2.0E-01 | 0.00 | 6.2E-01 | 0.05 | 4.8E-01 | -0.01 | 8.1E-01 | -0.01 | 9.3E-01 | 0.01 | 7.7E-01 | 0.00 | 9.7E-01 | 0.62 |
| FSIQ2 | 1.47 | 1.7E-01 | -25 | 0.01 | **5.8E-05** | -0.01 | **2.3E-02** | -0.02 | **7.9E-03** | 0.02 | **3.7E-02** | 1.54 | **2.2E-02** | 1.97 | **1.0E-02** | -2.07 | **4.3E-02** | 0.01 | 4.0E-01 | -0.01 | 8.5E-01 | **0.01** |
| PRI2 | 1.78 | 8.3E-02 | -28 | 0.01 | **4.8E-05** | 0.00 | 5.8E-01 | -0.02 | **3.8E-06** | 0.01 | 1.1E-01 | 0.35 | 5.9E-01 | 1.70 | **8.4E-06** | -1.23 | 1.4E-01 | 0.01 | 4.1E-01 | -0.01 | 8.5E-01 | **0.00** |
| PSI2 | 0.45 | 9.0E-01 | -16 | 0.01 | 1.6E-01 | -0.01 | 1.5E-01 | 0.00 | 3.5E-01 | 0.01 | 3.6E-01 | 0.75 | 1.2E-01 | 0.49 | 3.9E-01 | -0.61 | 3.8E-01 | 0.01 | 5.5E-01 | -0.01 | 7.7E-01 | 0.48 |

| | | | | | | | | | | | | | | | | | | | | | |
|---|---|---|---|---|---|---|---|---|---|---|---|---|---|---|---|---|---|---|---|---|---|
| VCI2 | 0.59 | 8.0E-01 | -17 | 0.00 | 5.6E-01 | -0.01 | 2.1E-01 | -0.01 | 4.2E-01 | 0.01 | 2.4E-01 | 0.92 | 2.0E-01 | 0.53 | 4.4E-01 | -1.07 | 2.5E-01 | 0.01 | 7.2E-01 | 0.00 | 9.6E-01 | 0.42 |
| WMI2 | 1.39 | 2.0E-01 | -25 | 0.01 | 1.7E-01 | -0.01 | 5.2E-01 | -0.01 | **3.6E-02** | 0.01 | 4.5E-01 | 0.54 | 5.1E-01 | 1.48 | **4.4E-02** | -0.66 | 4.8E-01 | 0.02 | 3.5E-01 | -0.01 | 8.3E-01 | **0.02** |
| Shen et al., (2015) | | | | | | | | | | | | | | | | | | | | | | |
| deltaFSIQ | 0.76 | 6.5E-01 | -7 | 0.01 | **3.7E-03** | -0.01 | 2.1E-01 | -0.02 | **3.3E-03** | 0.02 | 9.9E-02 | 0.07 | 3.6E-01 | 0.08 | 3.2E-01 | -0.09 | 3.4E-01 | 0.02 | 3.9E-01 | -0.01 | 8.3E-01 | 0.14 |
| deltaPRI | 1.77 | 8.6E-02 | -16 | 0.01 | 8.7E-02 | 0.00 | 8.4E-01 | -0.02 | **1.1E-03** | 0.01 | 1.2E-01 | 0.00 | 1.0E+00 | 0.06 | 3.5E-01 | -0.05 | 5.9E-01 | 0.02 | 2.8E-01 | -0.02 | 5.9E-01 | 0.00 |
| deltaPSI | 0.35 | 9.5E-01 | -3 | 0.00 | 2.8E-01 | 0.00 | 7.9E-01 | 0.00 | 5.8E-01 | 0.00 | 8.6E-01 | 0.04 | 6.1E-01 | -0.01 | 9.5E-01 | 0.01 | 9.3E-01 | 0.01 | 5.9E-01 | -0.01 | 8.1E-01 | 0.60 |
| deltaVCI | 0.64 | 7.6E-01 | -6 | 0.00 | 6.5E-01 | -0.01 | 1.1E-01 | 0.00 | 8.9E-01 | 0.01 | 4.1E-01 | 0.05 | 4.9E-01 | -0.02 | 7.2E-01 | -0.03 | 7.0E-01 | 0.01 | 6.5E-01 | -0.02 | 7.1E-01 | 0.22 |
| deltaWMI | 0.32 | 9.7E-01 | -3 | 0.00 | 5.4E-01 | 0.00 | 6.3E-01 | -0.01 | 2.4E-01 | 0.00 | 6.4E-01 | 0.04 | 5.5E-01 | -0.02 | 7.9E-01 | -0.01 | 8.8E-01 | 0.00 | 8.3E-01 | 0.00 | 9.4E-01 | 0.75 |
| FSIQ2 | 1.50 | 1.6E-01 | -14 | 0.01 | **2.4E-04** | -0.02 | **2.3E-02** | -0.02 | **1.9E-02** | 0.02 | 5.3E-02 | 1.68 | **2.3E-02** | 2.08 | **2.4E-02** | -2.20 | 5.9E-02 | 0.01 | 5.2E-01 | -0.01 | 8.7E-01 | 0.01 |
| PRI2 | 1.87 | 6.7E-02 | -17 | 0.01 | **9.7E-06** | 0.00 | 5.0E-01 | -0.02 | **3.2E-05** | 0.01 | 1.2E-01 | 0.47 | 5.1E-01 | 1.86 | **6.6E-05** | -1.35 | 1.4E-01 | 0.01 | 5.1E-01 | -0.01 | 8.3E-01 | 0.00 |
| PSI2 | 0.34 | 9.6E-01 | -3 | 0.00 | 2.3E-01 | -0.01 | 2.3E-01 | 0.00 | 4.6E-01 | 0.00 | 5.1E-01 | 0.67 | 1.9E-01 | 0.43 | 5.1E-01 | -0.48 | 5.2E-01 | 0.01 | 6.4E-01 | -0.01 | 7.9E-01 | 0.60 |
| VCI2 | 0.65 | 7.5E-01 | -6 | 0.00 | 5.6E-01 | -0.01 | 1.7E-01 | -0.01 | 4.6E-01 | 0.01 | 2.5E-01 | 1.06 | 1.7E-01 | 0.54 | 4.8E-01 | -1.15 | 2.5E-01 | 0.00 | 8.3E-01 | 0.00 | 9.7E-01 | 0.34 |
| WMI2 | 1.34 | 2.3E-01 | -13 | 0.01 | 1.4E-01 | -0.01 | 4.9E-01 | -0.02 | **2.5E-02** | 0.01 | 4.0E-01 | 0.58 | 4.8E-01 | 1.58 | **3.2E-02** | -0.77 | 4.2E-01 | 0.02 | 4.3E-01 | -0.01 | 8.4E-01 | 0.02 |
| Physics knowledge | | | | | | | | | | | | | | | | | | | | | | |

| | | | | | | | | | | | | | | | | | | | | | |
|---|---|---|---|---|---|---|---|---|---|---|---|---|---|---|---|---|---|---|---|---|---|
| Craddock et al., (2012) | | | | | | | | | | | | | | | | | | | | | |
| deltaFSIQ | 0.96 | 4.8E-01 | -2 | 0.01 | **4.2E-02** | 0.00 | 5.0E-01 | -0.02 | **8.4E-03** | 0.02 | 1.1E-01 | 0.02 | 7.5E-01 | 0.10 | 2.5E-01 | -0.08 | 4.4E-01 | 0.01 | 5.8E-01 | 0.03 | 4.9E-01 | 0.11 |
| deltaPRI | 1.85 | 6.9E-02 | -10 | 0.00 | 3.8E-01 | 0.00 | 7.0E-01 | -0.02 | **2.2E-03** | 0.01 | 7.6E-02 | 0.00 | 9.8E-01 | 0.07 | 3.1E-01 | -0.05 | 6.0E-01 | 0.01 | 7.8E-01 | 0.02 | 5.7E-01 | 0.00 |
| deltaPSI | 0.81 | 6.1E-01 | -1 | 0.00 | 5.3E-01 | 0.00 | 6.7E-01 | 0.00 | 3.9E-01 | 0.00 | 4.6E-01 | 0.00 | 9.8E-01 | 0.03 | 7.8E-01 | 0.06 | 6.2E-01 | 0.00 | 8.8E-01 | 0.03 | 4.0E-01 | 0.14 |
| deltaVCI | 0.77 | 6.4E-01 | -1 | 0.00 | 7.4E-01 | -0.01 | 3.9E-01 | 0.00 | 5.9E-01 | 0.01 | 5.9E-01 | 0.04 | 6.4E-01 | -0.03 | 6.7E-01 | -0.01 | 8.8E-01 | 0.00 | 9.2E-01 | 0.02 | 5.5E-01 | 0.25 |
| deltaWMI | 0.36 | 9.5E-01 | 3 | 0.01 | 3.1E-01 | 0.00 | 5.6E-01 | -0.01 | 3.6E-01 | 0.01 | 4.5E-01 | 0.03 | 6.6E-01 | 0.00 | 1.0E+00 | -0.02 | 8.5E-01 | 0.00 | 9.4E-01 | 0.03 | 4.3E-01 | 0.80 |
| FSIQ2 | 1.42 | 1.9E-01 | -7 | 0.01 | **3.0E-02** | -0.01 | 1.7E-01 | -0.02 | 2.7E-02 | 0.02 | 6.0E-02 | 1.14 | 1.6E-01 | 2.26 | **2.9E-02** | -2.27 | 6.5E-02 | 0.01 | 7.1E-01 | 0.03 | 4.9E-01 | 0.02 |
| PRI2 | 1.80 | 7.9E-02 | -10 | 0.01 | **5.6E-03** | 0.00 | 8.1E-01 | -0.02 | **6.1E-04** | 0.02 | 8.8E-02 | 0.17 | 8.3E-01 | 1.85 | **8.4E-04** | -1.61 | 1.1E-01 | 0.00 | 9.3E-01 | 0.03 | 3.9E-01 | 0.00 |
| PSI2 | 0.39 | 9.4E-01 | 3 | 0.00 | 3.3E-01 | 0.00 | 4.9E-01 | -0.01 | 3.2E-01 | 0.00 | 5.2E-01 | 0.42 | 4.4E-01 | 0.65 | 3.4E-01 | -0.49 | 5.3E-01 | 0.00 | 9.6E-01 | 0.03 | 4.2E-01 | 0.57 |
| VCI2 | 0.35 | 9.6E-01 | 3 | 0.00 | 5.4E-01 | -0.01 | 3.1E-01 | 0.00 | 5.6E-01 | 0.01 | 2.7E-01 | 0.67 | 2.9E-01 | 0.38 | 5.7E-01 | -0.93 | 2.7E-01 | 0.00 | 9.0E-01 | 0.03 | 4.4E-01 | 0.66 |
| WMI2 | 0.82 | 6.0E-01 | -1 | 0.01 | **2.3E-02** | -0.01 | 2.2E-01 | -0.01 | 1.7E-02 | 0.01 | 1.2E-01 | 0.88 | 2.0E-01 | 1.48 | **2.0E-02** | -1.35 | 1.3E-01 | 0.00 | 8.5E-01 | 0.03 | 4.7E-01 | 0.13 |
| Shen et al., (2015) | | | | | | | | | | | | | | | | | | | | | |
| deltaFSIQ | 0.89 | 5.4E-01 | 15 | 0.01 | **4.1E-02** | 0.00 | 5.3E-01 | -0.02 | **1.1E-02** | 0.02 | 1.3E-01 | 0.03 | 7.3E-01 | 0.10 | 2.9E-01 | -0.10 | 3.9E-01 | 0.01 | 5.3E-01 | 0.02 | 5.7E-01 | 0.15 |
| deltaPRI | 1.72 | 9.5E-02 | 7 | 0.00 | 4.0E-01 | 0.00 | 7.3E-01 | -0.02 | **5.0E-03** | 0.02 | 8.7E-02 | 0.00 | 9.7E-01 | 0.07 | 3.5E-01 | -0.07 | 5.1E-01 | 0.01 | 7.4E-01 | 0.02 | 6.3E-01 | 0.01 |

| | | | | | | | | | | | | | | | | | | | | | | |
|---|---|---|---|---|---|---|---|---|---|---|---|---|---|---|---|---|---|---|---|---|---|---|
| deltaPSI | 0.88 | 5.5E-01 | 15 | 0.00 | 4.9E-01 | 0.00 | 6.2E-01 | 0.00 | 3.7E-01 | -0.01 | 4.2E-01 | 0.00 | 9.6E-01 | 0.03 | 7.9E-01 | 0.05 | 6.5E-01 | 0.01 | 7.8E-01 | 0.03 | 4.8E-01 | 0.11 |
| deltaVCI | 0.78 | 6.4E-01 | 16 | 0.00 | 6.9E-01 | -0.01 | 3.6E-01 | 0.01 | 5.8E-01 | 0.01 | 6.1E-01 | 0.04 | 6.0E-01 | -0.03 | 6.5E-01 | -0.03 | 7.8E-01 | 0.00 | 9.9E-01 | 0.02 | 6.2E-01 | 0.26 |
| deltaWMI | 0.34 | 9.6E-01 | 20 | 0.01 | 3.2E-01 | 0.00 | 5.8E-01 | -0.01 | 4.4E-01 | 0.01 | 4.9E-01 | 0.04 | 6.3E-01 | 0.00 | 9.9E-01 | -0.03 | 7.3E-01 | 0.00 | 9.7E-01 | 0.03 | 5.0E-01 | 0.86 |
| FSIQ2 | 1.33 | 2.3E-01 | 11 | 0.01 | **3.3E-02** | -0.01 | 1.6E-01 | -0.02 | **3.7E-02** | 0.02 | 6.9E-02 | 1.26 | 1.6E-01 | 2.39 | **4.1E-02** | -2.48 | 7.3E-02 | 0.01 | 6.9E-01 | 0.03 | 5.5E-01 | 0.02 |
| PRI2 | 1.68 | 1.1E-01 | 8 | 0.01 | **1.0E-02** | 0.00 | 7.6E-01 | -0.02 | 2.9E-03 | 0.02 | 9.9E-02 | 0.26 | 7.7E-01 | 1.96 | **3.9E-03** | -1.83 | 1.2E-01 | 0.00 | 8.7E-01 | 0.03 | 4.6E-01 | 0.01 |
| PSI2 | 0.41 | 9.3E-01 | 19 | 0.00 | 3.0E-01 | 0.00 | 5.0E-01 | -0.01 | 2.9E-01 | 0.00 | 5.5E-01 | 0.43 | 4.5E-01 | 0.73 | 3.1E-01 | -0.51 | 5.5E-01 | 0.00 | 9.4E-01 | 0.03 | 4.8E-01 | 0.52 |
| VCI2 | 0.37 | 9.5E-01 | 20 | 0.00 | 5.2E-01 | -0.01 | 2.5E-01 | 0.00 | 5.7E-01 | 0.01 | 2.5E-01 | 0.79 | 2.3E-01 | 0.40 | 5.7E-01 | -1.05 | 2.4E-01 | 0.00 | 9.6E-01 | 0.03 | 5.0E-01 | 0.61 |
| WMI2 | 0.76 | 6.6E-01 | 16 | 0.01 | **2.4E-02** | -0.01 | 2.2E-01 | -0.02 | **2.3E-02** | 0.01 | 1.2E-01 | 0.96 | 2.0E-01 | 1.54 | 2.7E-02 | -1.48 | 1.3E-01 | 0.00 | 8.3E-01 | 0.03 | 5.2E-01 | 0.16 |

re, i.e., that the model is significant at $p_{FWE\text{-}corr} < 0.05$ and the WAIS-IV score regression coefficient is significant at $p < 0.05$. Bold model F-statistics indicate that the model is significant at $p_{FWE\text{-}corr} < 0.05$. Bold p-values indicate significant regression coefficients of significant relationships at $p < 0.05$. All model fit statistics (i.e., P(F)) were assessed for significance at a FWE-corrected $\alpha < 0.05$.

**Supplementary Table 5.** Significance of maximum t values of permuted ordinary least squares regressions of nodal efficiency on WAIS-IV scores.

| WAIS-IV Score | Regressor | Craddock et al., (2012) p(max t) | Shen et al., (2015) p(max t) |
|---|---|---|---|
| Physics Reasoning | | | |
| deltaPRI | WAIS | 4.24E-01 | 3.67E-01 |
| | WAISXSex | 4.24E-01 | 3.67E-01 |
| | WAISXClass | **6.40E-03** | 3.85E-02 |
| | WAISXSexXClass | 1.13E-01 | 5.34E-01 |
| | SexXClass | 6.01E-01 | 9.84E-01 |
| | Sex (F) | 6.60E-01 | 8.58E-01 |
| | Class (A) | 2.71E-01 | 5.92E-01 |
| | Age | 4.76E-01 | 8.54E-02 |
| | Year in Univ. | 4.16E-01 | 4.67E-01 |
| | **FD** | **3.00E-04** | **5.00E-04** |
| deltaFSIQ | WAIS | 3.00E-01 | 8.36E-01 |

| | | | |
|---|---|---|---|
| | WAISXSex | 3.00E-01 | 8.36E-01 |
| | WAISXClass | 1.97E-01 | 1.53E-01 |
| | WAISXSexXClass | 2.45E-01 | 6.37E-01 |
| | SexXClass | 3.39E-01 | 5.54E-01 |
| | Sex (F) | 9.51E-02 | 5.90E-01 |
| | Class (A) | 6.55E-01 | 6.21E-01 |
| | Age | 6.91E-01 | 1.26E-01 |
| | Year in Univ. | 2.98E-01 | 8.76E-01 |
| | **FD** | **1.00E-04** | **1.00E-04** |
| PRI2 | WAIS | 3.34E-01 | 6.28E-01 |
| | WAISXSex | 3.34E-01 | 6.28E-01 |
| | WAISXClass | 8.22E-02 | 9.10E-02 |
| | WAISXSexXClass | 2.74E-01 | 4.35E-01 |
| | SexXClass | 3.37E-01 | 3.74E-01 |
| | Sex (F) | 3.99E-01 | 6.48E-01 |

| | | | |
|---|---|---|---|
| | Class (A) | 1.08E-01 | 1.07E-01 |
| | Age | 7.56E-01 | 2.03E-01 |
| | Year in Univ. | 2.95E-01 | 9.52E-01 |
| | FD | 1.30E-03 | 1.00E-03 |
| FSIQ2 | WAIS | 3.53E-02 | 6.20E-02 |
| | WAISXSex | 3.53E-02 | 6.20E-02 |
| | WAISXClass | 4.54E-01 | 1.20E-01 |
| | WAISXSexXClass | 1.15E-01 | 1.90E-01 |
| | SexXClass | 1.21E-01 | 2.14E-01 |
| | Sex (F) | 3.78E-02 | 5.76E-02 |
| | Class (A) | 4.50E-01 | 1.09E-01 |
| | Age | 5.44E-01 | 2.64E-01 |
| | Year in Univ. | 2.70E-01 | 9.76E-01 |
| | **FD** | **2.00E-04** | **4.00E-04** |
| Physics knowledge | | | |

| | | | |
|---|---|---|---|
| WMI2 | WAIS | 2.45E-01 | 8.23E-01 |
| | WAISXSex | 2.45E-01 | 8.23E-01 |
| | WAISXClass | 7.99E-01 | 9.59E-01 |
| | WAISXSexXClass | 2.02E-02 | 3.94E-01 |
| | SexXClass | 2.45E-02 | 3.76E-01 |
| | Sex (F) | 2.22E-01 | 7.92E-01 |
| | Class (A) | 8.21E-01 | 9.09E-01 |
| | Age | 4.92E-01 | 9.67E-01 |
| | Year in Univ. | 5.95E-01 | 6.66E-01 |
| | FD | 2.31E-02 | 1.91E-02 |
| VCI2 | WAIS | 2.15E-01 | 5.44E-01 |
| | WAISXSex | 2.15E-01 | 5.44E-01 |
| | WAISXClass | 3.04E-01 | 4.71E-02 |
| | WAISXSexXClass | 2.84E-01 | 5.24E-01 |
| | SexXClass | 2.89E-01 | 4.25E-01 |

| | | |
|---|---|---|
| Sex (F) | 1.96E-01 | 5.16E-01 |
| Class (A) | 3.28E-01 | 4.54E-02 |
| Age | 3.97E-01 | 9.26E-01 |
| Year in Univ. | 4.34E-01 | 6.74E-01 |
| FD | 1.01E-02 | 1.86E-02 |

Bold p-values indicate parameters for which at least one node's efficiency was significantly related to a regressor at $p_{FWE\text{-}corr} < 0.05$. Bold regressors indicate that at least one node's efficiency was significantly related to the regressor in both parcellations.

**Supplementary Table 6**. Significance of maximum t values from permuted ordinary least squares regression of connectivity on WAIS scores.

| WAIS-IV Score | Regressor | Shen et al., (2015) | Craddock et al., (2012) |
|---|---|---|---|
| Physics reasoning | | | |
| PRI2 | WAIS | 2.58E-01 | 3.48E-01 |
| | WAISXSex | 9.46E-02 | 1.61E-01 |
| | WAISXClass | 4.31E-01 | 5.20E-01 |
| | WAISXSexXClass | 2.59E-02 | 4.06E-02 |
| | SexXClass | 1.88E-02 | 3.48E-02 |
| | Sex (F) | 9.92E-02 | 1.68E-01 |
| | Class (A) | 4.73E-01 | 4.45E-01 |
| | Age | 3.30E-01 | 8.28E-02 |
| | Year in Univ | **4.80E-03** | 5.94E-02 |
| | FD | 1.08E-02 | 2.27E-02 |
| **FSIQ2** | WAIS | 3.65E-02 | 1.42E-02 |

| | | | |
|---|---|---|---|
| | WAISXSex | 1.47E-02 | **4.90E-03** |
| | WAISXClass | 3.34E-02 | **1.20E-03** |
| | **WAISXSexXClass** | **1.70E-03** | **2.50E-03** |
| | **SexXClass** | **1.80E-03** | **2.70E-03** |
| | Sex (F) | 1.25E-02 | **4.20E-03** |
| | Class (A) | 3.15E-02 | **1.60E-03** |
| | Age | 5.19E-01 | 1.64E-01 |
| | Year in Univ | **4.00E-03** | 6.36E-02 |
| | FD | **4.50E-03** | 2.22E-02 |
| deltaPRI | WAIS | 6.71E-02 | 7.61E-02 |
| | WAISXSex | 1.62E-01 | 1.24E-01 |
| | WAISXClass | 1.91E-01 | 1.99E-01 |
| | WAISXSexXClass | 1.56E-02 | **4.20E-03** |
| | SexXClass | 8.60E-03 | 3.80E-02 |
| | Sex (F) | 5.47E-02 | **5.20E-03** |

| | | | |
|---|---|---|---|
| | Class (A) | 6.02E-01 | 2.81E-01 |
| | Age | 4.98E-01 | 1.63E-01 |
| | Year in Univ | 1.88E-02 | 1.44E-01 |
| | FD | 9.10E-03 | 1.20E-02 |
| deltaFSIQ | WAIS | 4.25E-02 | 2.41E-02 |
| | WAISXSex | 2.38E-02 | 2.45E-02 |
| | WAISXClass | 2.09E-01 | 4.56E-01 |
| | WAISXSexXClass | 3.51E-02 | 4.15E-02 |
| | SexXClass | 1.90E-01 | 4.12E-02 |
| | Sex (F) | **4.80E-03** | 1.20E-01 |
| | Class (A) | 5.83E-01 | 1.76E-01 |
| | Age | 6.61E-01 | 5.08E-01 |
| | Year in Univ | 1.05E-02 | 1.77E-02 |
| | FD | 1.20E-02 | 2.52E-02 |
| Physics knowledge | | | |

| | | | |
|---|---|---|---|
| VCI2 | WAIS | 1.60E-01 | 1.30E-01 |
| | WAISXSex | 2.61E-01 | 3.87E-01 |
| | WAISXClass | 4.59E-02 | 4.67E-01 |
| | WAISXSexXClass | 5.52E-01 | 6.54E-01 |
| | SexXClass | 5.07E-01 | 6.08E-01 |
| | Sex (F) | 2.34E-01 | 4.44E-01 |
| | Class (A) | 4.69E-02 | 5.37E-01 |
| | Age | 1.69E-01 | 1.10E-01 |
| | Year in Univ | 4.32E-02 | 3.86E-02 |
| | **FD** | **4.80E-03** | **3.00E-04** |
| WMI2 | WAIS | 7.18E-02 | 1.20E-01 |
| | WAISXSex | 1.36E-01 | 1.66E-01 |
| | WAISXClass | 1.04E-01 | 9.44E-02 |
| | WAISXSexXClass | 6.88E-02 | 4.85E-02 |
| | SexXClass | 6.67E-02 | 5.89E-02 |

| | | |
|---|---|---|
| Sex (F) | 1.40E-01 | 2.07E-01 |
| Class (A) | 1.12E-01 | 9.31E-02 |
| Age | 5.94E-01 | 3.02E-01 |
| Year in Univ | 3.45E-02 | 3.97E-02 |
| FD | 1.24E-02 | **1.60E-03** |

Bold p-values indicate parameters for which at least one node's efficiency was significantly related to a regressor at $p_{FWE\text{-}corr} < 0.05$. Bold regressors indicate that at least one node's efficiency was significantly related to the regressor in both parcellations. Bold WAIS-IV scores indicate a regression in which a regressor including that WAIS score was significant in both parcellations.

**Supplementary Table 7**. Results of Maximum Likelihood estimated models of task-based connectivity mediating the relationship between WAIS scores and physics reasoning accuracy

| | Craddock et al., (2012) | | | Shen et al., (2015) | | |
|---|---|---|---|---|---|---|
| | Estimate | z-value | P(Estimate) | Estimate | z-value | P(Estimate) |
| Model test statistic ( $\chi^2$) | 262.216 | -- | **0.000** | 262.848 | -- | **0.000** |
| RMSEA | 0.419 | | **0.000** | 0.420 | | **0.000** |
| Edge ~ | $R^2 = 0.154$ | | | $R^2 = 0.157$ | | |
| FSIQ2 | -0.009 | -0.392 | 0.695 | 0.03 | 1.205 | 0.228 |
| FSIQXSex | 0.062 | 2.247 | 0.025 | 0.048 | 1.411 | 0.158 |
| FSIQXClass | 0.004 | 0.134 | 0.893 | -0.024 | -0.631 | 0.528 |
| FSIQXSexXClass | -0.071 | -2.026 | 0.043 | -0.072 | -1.211 | 0.226 |
| Age | -0.001 | -0.086 | 0.931 | -0.003 | -0.267 | 0.789 |
| Year in Univ. | 0.002 | 0.727 | 0.467 | 0.004 | 1.848 | 0.065 |

| | | | | | | |
|---|---|---|---|---|---|---|
| SexXClass | 0.046 | 1.286 | 0.199 | -0.028 | -0.553 | 0.58 |
| Sex (F) | -0.037 | -1.126 | 0.260 | 0.01 | 0.359 | 0.72 |
| Class (A) | -0.033 | -1.426 | 0.154 | -0.003 | -0.064 | 0.949 |
| **FD** | 0.021 | 1.654 | 0.098 | 0.032 | 2.028 | **0.043** |
| Physics reasoning accuracy ~ $R^2 = 0.240$ | | | | $R^2 = 0.244$ | | |
| FSIQ2 | 0.054 | 1.543 | 0.123 | 0.056 | 1.603 | 0.109 |
| FSIQXSex | 0.011 | 0.247 | 0.805 | 0.015 | 0.35 | 0.727 |
| FSIQXClass | 0.003 | 0.078 | 0.938 | 0.001 | 0.03 | 0.976 |
| FSIQXSexXClass | 0.023 | 0.398 | 0.691 | 0.015 | 0.252 | 0.801 |
| Age | -0.013 | -1.38 | 0.168 | -0.014 | -1.39 | 0.165 |
| Year in Univ. | 0 | 0.041 | 0.967 | 0 | 0.178 | 0.859 |
| SexXClass | 0.053 | 0.926 | 0.355 | 0.051 | 0.876 | 0.381 |
| **Sex (F)** | -0.135 | -3.379 | **0.001** | -0.134 | -3.304 | **0.001** |

| | | | | | | |
|---|---|---|---|---|---|---|
| Class (A) | -0.056 | -1.308 | 0.191 | -0.057 | -1.328 | 0.184 |
| Edge | 0.02 | 0.137 | 0.891 | -0.071 | -0.912 | 0.362 |
| Covariances | | | | | | |
| FSIQ2XSex | ~~ | | | ~~ | | |
| **FSIQ2XClass** | 0.217 | 2.614 | **0.009** | 0.217 | 2.614 | **0.009** |
| **FSIQ2XClassXSex** | 0.203 | 2.526 | **0.012** | 0.203 | 2.526 | **0.012** |
| Age | -0.139 | -1.515 | 0.13 | -0.139 | -1.515 | 0.13 |
| Year in Univ. | -0.249 | -0.585 | 0.559 | -0.249 | -0.585 | 0.559 |
| SexXClass | -0.053 | -1.657 | 0.097 | -0.053 | -1.657 | 0.097 |
| Sex (F) | -0.08 | -2.729 | **0.006** | -0.08 | -2.729 | **0.006** |
| Class (A) | -0.009 | -0.322 | 0.747 | -0.009 | -0.322 | 0.747 |
| fd | -0.113 | -0.793 | 0.428 | -0.113 | -0.793 | 0.428 |
| FSIQ2XClass | ~~ | | | ~~ | | |
| **FSIQ2XClassXSex** | 0.216 | 2.547 | **0.011** | 0.216 | 2.547 | **0.011** |

| | | | | | | |
|---|---|---|---|---|---|---|
| Age | -0.062 | -0.559 | 0.576 | -0.062 | -0.559 | 0.576 |
| Year in Univ. | 0.096 | 0.163 | 0.871 | 0.096 | 0.163 | 0.871 |
| **SexXClass** | -0.085 | -2.525 | **0.012** | -0.085 | -2.525 | **0.012** |
| **Sex (F)** | -0.088 | -2.738 | **0.006** | -0.088 | -2.738 | **0.006** |
| Class (A) | 0.006 | 0.188 | 0.851 | 0.006 | 0.188 | 0.851 |
| fd | -0.153 | -0.984 | 0.325 | -0.153 | -0.984 | 0.325 |
| FSIQ2XClassXSex | ~~ | | | ~~ | | |
| Age | -0.016 | -0.425 | 0.671 | -0.016 | -0.425 | 0.671 |
| Year in Univ. | 0.021 | 0.096 | 0.924 | 0.021 | 0.096 | 0.923 |
| **SexXClass** | -0.066 | -2.117 | **0.034** | -0.066 | -2.117 | **0.034** |
| **Sex (F)** | -0.044 | -2.096 | **0.036** | -0.044 | -2.096 | **0.036** |
| **Class (A)** | -0.042 | -2.092 | **0.036** | -0.042 | -2.092 | **0.036** |
| fd | -0.147 | -1.027 | 0.305 | -0.147 | -1.027 | 0.305 |
| Age | ~~ | | | ~~ | | |
| **Year in Univ.** | 5.419 | 4.709 | **0** | 5.419 | 4.709 | **0** |

| | | | | | | |
|---|---|---|---|---|---|---|
| SexXClass | 0.041 | 0.874 | 0.382 | 0.041 | 0.874 | 0.382 |
| Sex (F) | 0.023 | 0.387 | 0.698 | 0.023 | 0.387 | 0.698 |
| Class (A) | 0.013 | 0.222 | 0.824 | 0.013 | 0.222 | 0.824 |
| fd | -0.059 | -0.562 | 0.574 | -0.059 | -0.562 | 0.574 |
| Year in Univ. | ~~ | | | ~~ | | |
| SexXClass | 0.215 | 0.814 | 0.416 | 0.215 | 0.814 | 0.416 |
| Sex (F) | 0.784 | 2.303 | 0.021 | 0.784 | 2.303 | 0.021 |
| Class (A) | -0.174 | -0.503 | 0.615 | -0.174 | -0.503 | 0.615 |
| fd | -0.544 | -0.986 | 0.324 | -0.544 | -0.986 | 0.324 |
| SexXClass | ~~ | | | ~~ | | |
| **Sex (F)** | 0.108 | 6.745 | **0** | 0.108 | 6.745 | **0** |
| **Class (A)** | 0.102 | 6.616 | **0** | 0.102 | 6.616 | **0** |
| fd | 0.007 | 0.148 | 0.882 | 0.007 | 0.148 | 0.882 |
| Sex (F) | ~~ | | | ~~ | | |
| Class (A) | -0.027 | -1.253 | 0.21 | -0.027 | -1.253 | 0.21 |

| | | | | | | |
|---|---|---|---|---|---|---|
| fd | -0.003 | -0.063 | 0.949 | -0.003 | -0.063 | 0.949 |
| Class (A) | ~~ | | | ~~ | | |
| fd | 0.013 | 0.287 | 0.774 | 0.013 | 0.287 | 0.774 |
| Defined parameters | | | | | | |
| Simple slopes | | | | | | |
| WAIS - Female | 0.115 | 2.903 | 0.004 | 0.126 | 2.352 | **0.019** |
| WAIS - Male | 0.053 | 3.081 | 0.002 | 0.078 | 3.185 | **0.001** |
| WAIS - Active | -0.002 | -0.039 | 0.969 | -0.018 | -0.28 | 0.78 |
| WAIS - Lecture | -0.005 | -0.351 | 0.725 | 0.006 | 0.198 | 0.843 |
| WAIS - Active, Female | 0.122 | 1.517 | 0.129 | 0.077 | 0.73 | 0.465 |
| WAIS - Lecture, Female | 0.118 | 2.088 | 0.037 | 0.101 | 1.361 | 0.174 |
| WAIS - Active, Male | 0.06 | 1.058 | 0.29 | 0.03 | 0.365 | 0.715 |
| WAIS - Lecture, Male | 0.057 | 1.821 | 0.069 | 0.054 | 1.171 | 0.242 |
| Conditional indirect effects | | | | | | |
| WAISxConn Female | 0.002 | 0.136 | 0.891 | -0.009 | -0.861 | 0.389 |

| | | | | | | |
|---|---|---|---|---|---|---|
| WAISXConn Male | 0.001 | 0.137 | 0.891 | -0.006 | -0.878 | 0.38 |
| WAISXConn Active | 0 | -0.038 | 0.969 | 0.001 | 0.268 | 0.789 |
| WAISXConn Lecture | 0 | -0.129 | 0.897 | 0 | -0.192 | 0.848 |
| WAISXConn Active, Female | 0.002 | 0.135 | 0.893 | -0.005 | -0.579 | 0.563 |
| WAISXConn Lecture, Female | 0.002 | 0.136 | 0.892 | -0.007 | -0.77 | 0.441 |
| WAISXConn Lecture, Male | 0.001 | 0.136 | 0.892 | -0.004 | -0.726 | 0.468 |
| WAISXConn Active, Male | 0.001 | 0.134 | 0.893 | -0.002 | -0.341 | 0.733 |
| Indices of moderated mediation | | | | | | |
| modMedSex | 0.001 | 0.136 | 0.892 | -0.003 | -0.786 | 0.432 |
| modMedClass | 0 | 0.09 | 0.928 | 0.002 | 0.515 | 0.607 |
| modMedSexClass | -0.001 | -0.136 | 0.892 | 0.005 | 0.723 | 0.469 |
| edgeMedIQAcc | 0 | -0.125 | 0.901 | -0.002 | -0.704 | 0.481 |

Bold values indicate significant parameter estimates. Bold regressors indicate regressors whose estimates are significant in mediations run with connectivity values from both parcellations.

### Supplemental Figures

**Supplementary Figure 1.** Distribution of pre- and post-instruction WAIS-IV scores and index scores across the sample, and pre-to-post changes in those scores

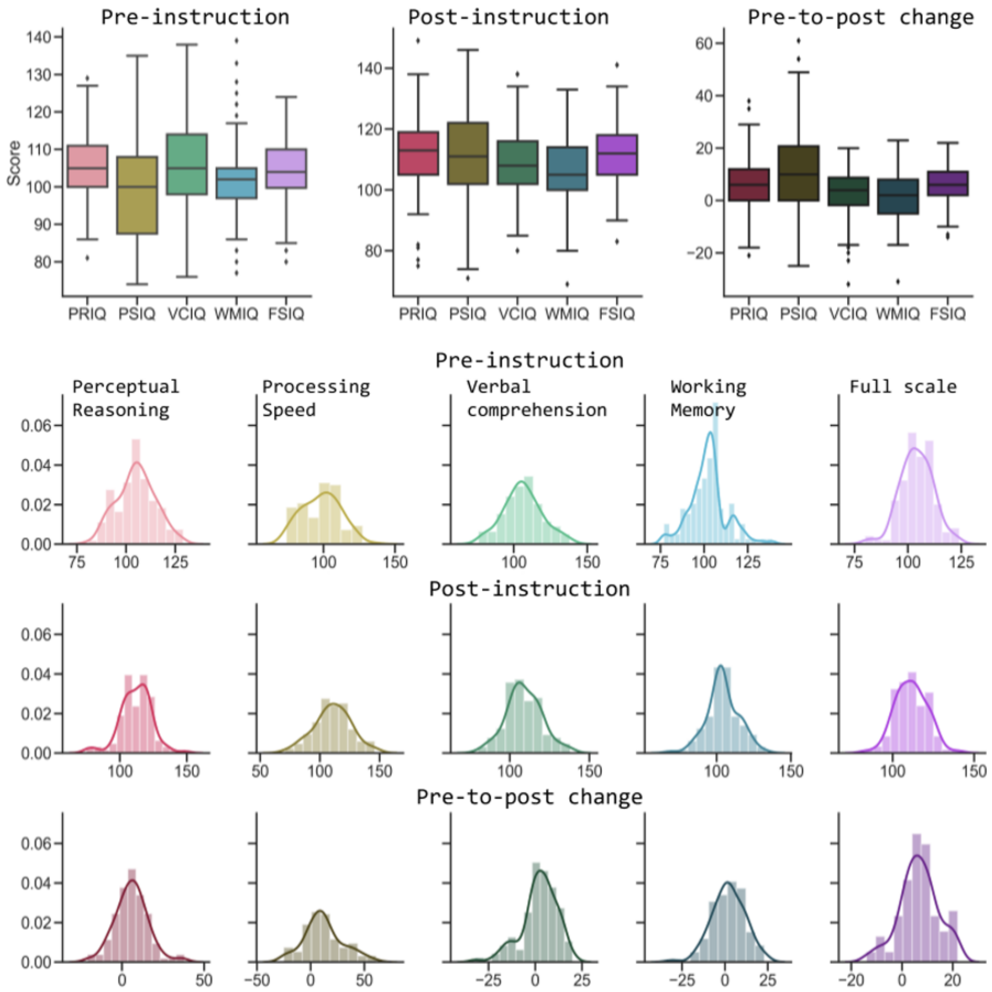


.

**Supplementary Figure 2.** Functional connectivity during physics reasoning and knowledge tasks in the Craddock Parcellation

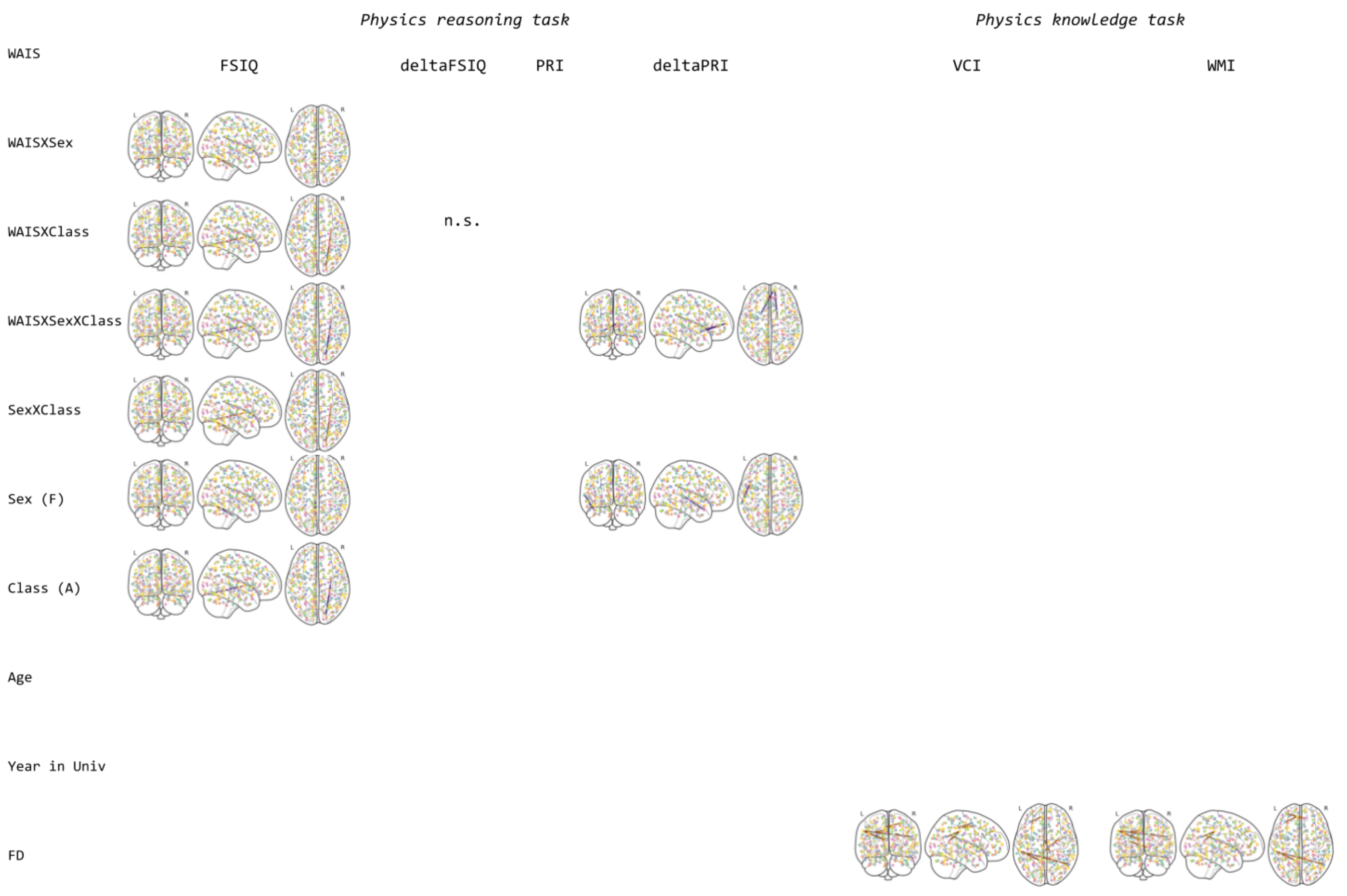

**Supplementary Figure 3.** Functional connectivity during physics reasoning and knowledge tasks in the Shen Parcellation

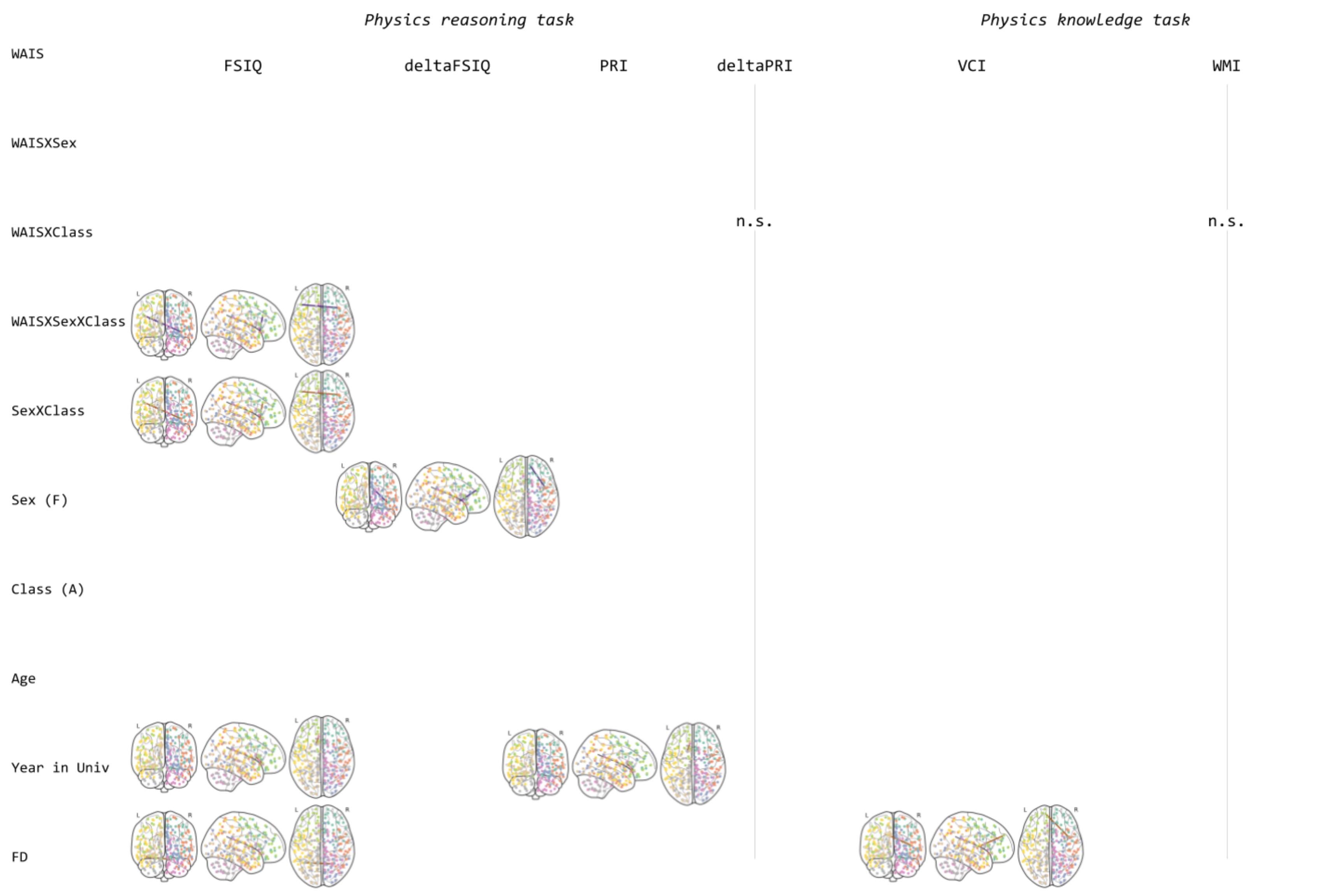

**Supplementary Figure 4.** Brain parcellations used to define nodes across which brain network connectivity was calculated.

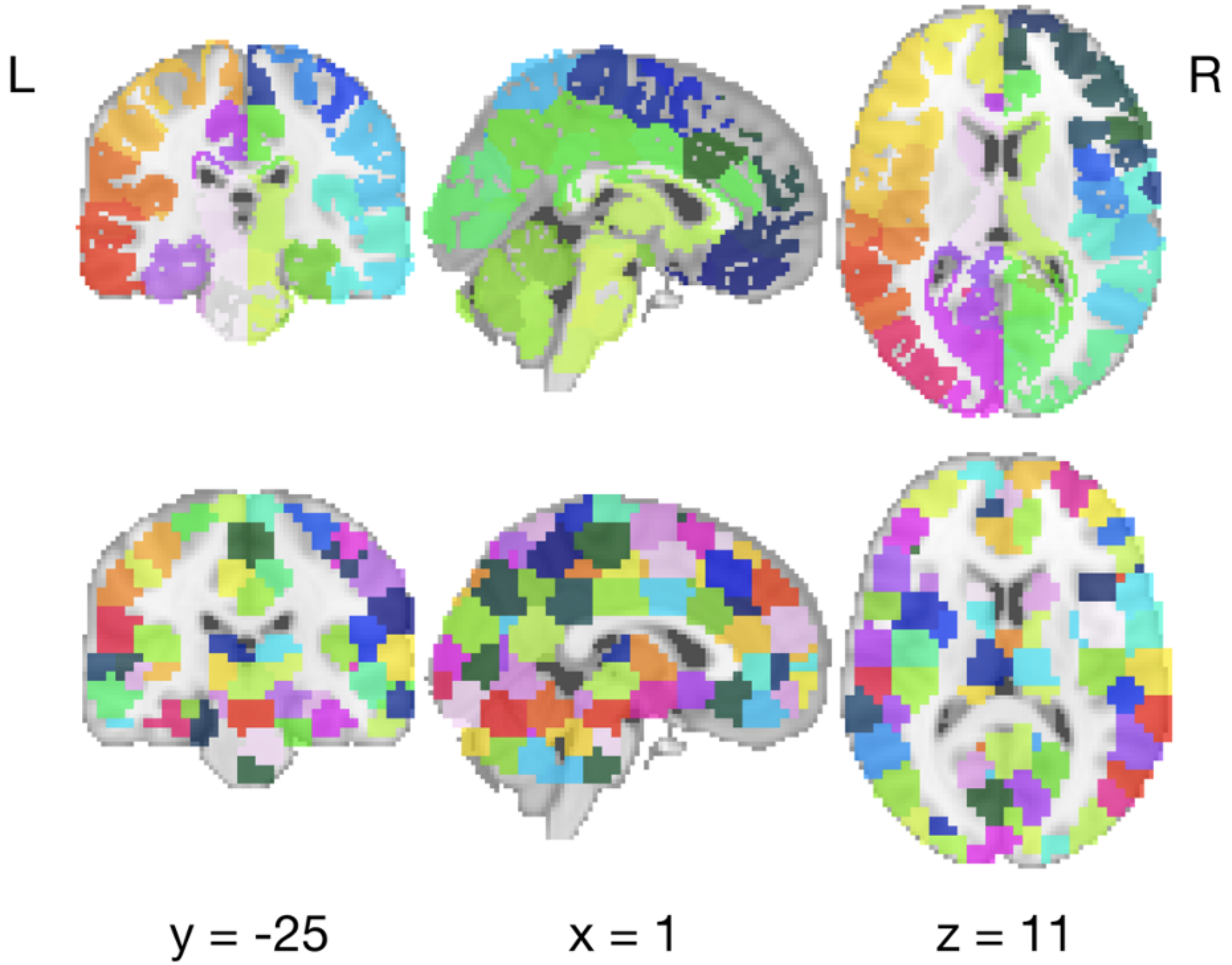


Top: 268-region parcellation computed via multigraph *k*-way clustering, without spatial constraints, by Shen et al., (2015). Bottom: 268-region parcellation computed via normalized-cut spectral clustering to define 268 homogenous, spatially-constrained clusters (i.e., regions) by Craddock et al. (2012).
